# A cerebellar-prepontine circuit for tonic immobility triggered by inescapable threat

**DOI:** 10.1101/2021.09.02.458709

**Authors:** Ashwin A Bhandiwad, Nickolas Chu, Svetlana A Semenova, Harold A Burgess

## Abstract

Sudden changes in the sensory environment are frequently perceived as threats and may provoke defensive behavioral states. One such state is tonic immobility, a conserved defensive strategy characterized by a powerful suppression of movement and motor reflexes. Tonic immobility has been associated with multiple brainstem regions and cell types, but the underlying circuit is not known. Here, we demonstrate that a strong vibratory stimulus evokes tonic immobility in larval zebrafish defined by suppression of exploratory locomotion and sensorimotor responses. Using a circuit-breaking screen and targeted neuron ablations, we show that cerebellar granule cells and a cluster of glutamatergic ventral prepontine neurons (vPPNs) that express key stress-associated neuropeptides are critical components of the circuit that suppresses movement. The complete sensorimotor circuit transmits information from primary sensory neurons through the cerebellum to vPPNs to regulate reticulospinal premotor neurons. These results show that cerebellar regulation of a neuropeptide-rich prepontine structure governs a conserved and ancestral defensive behavior that is triggered by inescapable threat.

## Introduction

Sudden changes in the environment may signal new threats. In response, animals often avoid detection by predators and facilitate threat assessment by suppressing physical activity (Klemm, 2001; Roseberry and Kreitzer, 2017; Yeomans and Frankland, 1995). Behavioral arrest is a common defensive strategy against predatory threat described in many vertebrate and invertebrate species (Gibson et al., 2015; Liang et al., 2015; Perrins et al., 2002; Sisneros et al., 1998; Zacarias et al., 2018) that forms part of the defense cascade, a continuum of behaviors that scale with perceived threat immediacy (Fanselow, 1994; Gallup, 1977; Kozlowska et al., 2015; Marx et al., 2008). Physical activity is suppressed at each end of the defense cascade as part of responses to both distant and immediate threat. When peril is relatively low, for example at first detection of a predator cue, animals instigate avoidance behaviors. In the post-encounter phase of defensive behaviors, animals exhibit ‘freezing’ behavior, characterized by suppressed movement and heightened sensory acuity with increased readiness to flee. With escalating threat, animals initiate escape behaviors, and finally, when faced with imminent entrapment or actual capture, lapse into a catatonic-like state of ‘tonic immobility’. The defense cascade presents a valuable model for resolving neural mechanisms that select and coordinate the expression of competing behavioral programs.

Freezing and tonic immobility are distinct behaviors within the defense cascade. Both behaviors are defined by behavioral arrest but are triggered by different levels of perceived threat and have converse effects on sensory responsiveness, muscle tone and postural control (Kozlowska et al., 2015). The core circuit for freezing behavior consists of the periaqueductal gray, although additional regions including the parabrachial nuclei, cortex, and cerebellum have been implicated (Dean et al., 1989; LeDoux and Daw, 2018; Liang et al., 2015; Roelofs, 2017; Roseberry and Kreitzer, 2017; Tovote et al., 2016; Vaaga et al., 2020). In contrast, very little is known about the neural circuit basis for tonic immobility. States of tonic immobility have been described under different names, including ‘phasic immobility’, ‘playing possum’, ‘death feigning’, and ‘thanatosis’ (Humphreys and Ruxton, 2018; Rogers and Simpson, 2014). In humans, tonic immobility is linked to feelings of paralysis during traumatic events and is a major predictor of post-traumatic stress disorder severity (Kalaf et al., 2015; Marx et al., 2008; Volchan et al., 2017). Imminent threat or restraint induces a state in which animals are motionless and unreactive to external stimuli, possibly to reduce immediate predator aggression with the goal of escaping later. Paradoxically, animals in this state show increased EEG theta power, heart rate, and breathing rate, suggesting hyperarousal (Klemm, 2001). Early decerebration studies showed that the circuit for restraint-induced tonic immobility was localized to the brainstem (McBride and Klemm 1969, Klemm 1977) but the specific neural substrates and circuit connectivity for inducing tonic immobility are still unknown. How extreme threats initiate tonic immobility, over-riding the expression of alternative defensive behaviors, remains a fundamental question.

Both freezing behavior and tonic immobility occur in fishes. Freezing behavior has been studied as a response to electrical shock (Agetsuma et al., 2010; Duboué et al., 2017), exposure to alarm pheromones (Jesuthasan et al., 2020; Maximino et al., 2019; Speedie and Gerlai, 2008), and placement into a novel tank (Cachat et al., 2010). Several studies have also reported a state of tonic immobility in fish that manifests as prolonged inactivity, loss of responsiveness and suppression of righting reflexes, often after manual restraint or physical inversion (Assad et al., 2020; Carli, 1968; Crawford, 1977; Henningsen, 1994; Lefebvre and Sabourin, 1977; Yoshida, 2021). In larval stage zebrafish, immobility is induced by an extreme vibratory stimulus well above intensity levels that normally induce escape responses raising the possibility of applying circuit neuroscience methods to unraveling a neuronal substrate for this behavior (Yokogawa et al., 2012).

Here, we demonstrate tonic immobility in larval zebrafish evoked by a persistent and inescapable threat and defined by behavioral arrest and reflex suppression. Using a genetic circuit breaking screen, targeted ablations, and imaging, we map a contiguous sensorimotor pathway for this terminal defensive behavior from primary mechanosensory neurons via the cerebellum and vPPNs to premotor reticulospinal neurons.

We show that vPPNs express multiple stress and homeostasis-related neuropeptides and are bilaterally activated by threatening stimuli. Finally, we propose a circuit model where multimodal sensory inputs act through the cerebellum to bilaterally activate vPPNs and trigger behavioral arrest by disrupting RoL1 reticulospinal neuron activity. Together, these results reveal a novel sensorimotor circuit for this highly conserved defensive behavior that is activated when faced with an inescapable threat.

## Results

### Repeated vibratory stimuli evoke locomotor arrest

Intense vibratory stimuli elicit a sustained decrease in locomotor activity in zebrafish larvae (Yokogawa et al., 2012). To characterize this state, we delivered repeated pulsed high amplitude, low frequency stimulus using a vibration exciter and measured changes in swim speed relative to a 60 second pre-stimulus baseline (Fig 1A-B). Individual pulses elicited escape responses with 83 ± 5% responsiveness and the stimulus was repeated for a 15 sec period – this simulated a persistent and inescapable threat. Following repeated stimulus presentation, larvae showed a persistent reduction in swimming which scaled with stimulus amplitude (Fig 1C-D). This reduction recovered to the pre-stimulus baseline level over the subsequent minute, with a half recovery time (T_0.5_) of 14-26 s (95% CI, n = 27 larvae). For all subsequent experiments, we used a stimulus that decreased speed by 50% with a half-recovery time of 20 s (Fig S1A).

**Figure 1:**
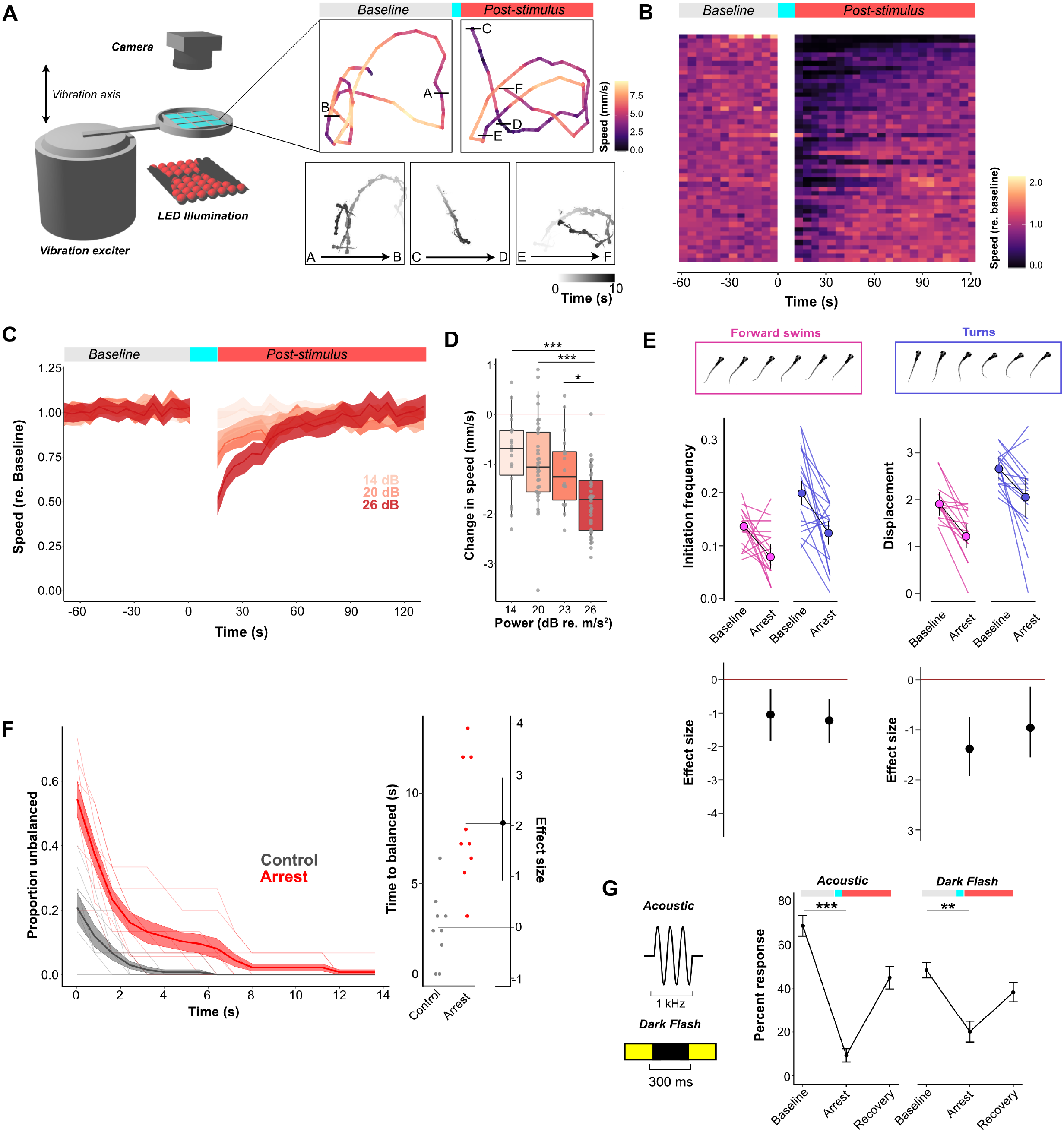
Repeated vibratory stimuli evoke tonic immobility with behavioral arrest and reflex suppression. A (Left) Schematic of apparatus used to evoke behavioral arrest. Individual fish are placed into a single well of a 3×3 grid plate attached to a minishaker that delivers vibratory stimuli. The arena is lit from below using an LED array and imaged using a low (30 fps) or high-speed (1000 fps) camera. (Right, Top) Individual locomotion traces from a single fish during baseline and post-stimulus phases. Color represents speed in a 5 sec epoch. (Right, Bottom) Individual fish positions during 10 s epochs of baseline (A, B) arrest (C, D) and recovery (E, F) phases – saturation indicates time within epoch. B Raster plot of individual fish speed (mm/s; n = 51 fish) for the duration of the experiment. Each line represents speed of an individual fish measured in 5 s epochs and color represents speed relative to the average baseline speed. Gap in data corresponds to vibratory stimulus presentation. C Arrest magnitude is proportional to stimulus magnitude. Traces showing speed (relative to the average baseline speed) after 15 s arrest-inducing stimuli of 0 dB (no stimulus), 14 dB, 20 dB, and 26 dB re. 1 m/s^2^ (n = 27 fish). Shaded area for each trace is SEM. Arrest magnitude is stimulus-dependent (ANOVA; p < 0.001). D Boxplots showing change in speed (mm/s) at the first epoch following stimulus presentation compared with the pre-stimulus baseline at 14 dB, 20 dB, 23 dB, and 26 dB re. 1 m/s^2^ (n = 27 fish per condition). * p < 0.05, *** p < 0.001 E Kinematic analysis of changes in initiation frequency and displacement during arrest for turns (blue) and forward swims (magenta). (Top) Examples of turns and forward swims. (Middle) Individual traces showing changes in initiation frequency (Left) and displacement for each bout (Right) during baseline and arrest states. (Bottom) Estimated effect sizes for difference in responsiveness after stimulus compared to pre-stimulus baseline. Data plotted as Mean ± 95% CI. Horizontal line denotes an effect size of 0. F (Left) Righting reflex, measured as the proportion of unbalanced fish following destabilizing stimulus (n = 10 groups with 5 fish per group) in control (grey) and arrest (red) conditions. Individual lines represent groups of 5 fish and shaded area is SEM. (Right) Estimation plot for time to return to balanced calculated for each group of fish in control (grey) and arrest (red) conditions. G (Left) schematic diagram of acoustic and dark flash stimuli. (Right) Changes in acoustic short-latency startle (n = 27 fish) and dark-flash evoked O-bend responses (n = 27 fish) during arrest (Mean ± SEM). Responsiveness is significantly suppressed during arrest compared to baseline ** p < 0.01, *** p < 0.001, Repeated-Measures ANOVA.

After the intense stimulus, remaining swim activity showed relatively straight trajectories compared to the looping paths observed during baseline conditions (Fig 1A, S1B). Kinematic analysis of high-speed video indicated that this was due to a change in swim bout type usage. Spontaneous movement in larvae primarily occurs as discrete bouts of slow forward swims and routine turns (Fero et al., 2011; Marques et al., 2018). Consistent with the switch to straight swim trajectories, during arrest, larvae showed a large reduction in the frequency of turn movements, with a smaller decrease in forward swim initiations (Fig 1E). Displacement per movement was also decreased (Fig 1E). Thus, during vibration-induced arrest, the decreased movement is due to a reduction in both initiation frequency of movement events and in the travel distance of each event.

In other species, reduced movement to threats can be classified as freezing or tonic immobility through differential effects on sensorimotor reflexes including modulation of the righting reflex, a vestibular reflex that restores dorsoventral orientation when the animal is restrained and inverted (Jänicke and Coper, 1996) — freezing facilitates whereas tonic immobility suppresses these responses (Fanselow, 1994; Gallup and Rager, 1996). Therefore, to differentiate between freezing and tonic immobility, we investigated changes to sensorimotor reflexes during behavioral arrest.

At baseline, zebrafish consistently exhibit a dorsal-up posture. As in other animals, inversion or forced side-lying evokes a vestibulomotor self-righting reflex to restore the dorsal-up posture (Bagnall and McLean, 2014; Favre-Bulle et al., 2017). We tested whether the righting reflex was suppressed during behavioral arrest by presenting a brief high intensity low frequency stimulus that disrupted balance. Destabilizing larvae during the arrest state increased the likelihood of inversion and delayed restoration of the dorsal-up posture compared to baseline, demonstrating a disruption of the righting reflex, consistent with tonic immobility (Fig 1F). Moreover, acoustic-evoked startle and visual reflexes were suppressed during arrest (Fig 1G). Thus, after intense vibration, zebrafish show arrested movement, suppression of sensorimotor responses and loss of the righting reflex, all characteristic features of tonic immobility as manifest in other species under extreme threat.

Early work in rabbits suggested that immobility reflected inhibition of spinal motor circuits (Klemm, 1976). We tested whether motor circuit function was altered during arrest by direct activation of Mauthner cells with an electrical pulse stimulus (Tabor et al., 2014). Electric pulse-initiated escapes were normal and kinematic analysis of remaining visual and acoustic responses performed during arrest showed no changes in escape bend angle or displacement (Fig S1C, 2). It is therefore unlikely that motor circuits are directly suppressed during behavioral arrest. Rather, reduced movement reflects a change in premotor activity that initiates and sustains movement.

To reveal neurons that mediate behavioral arrest, we conducted a circuit breaking screen using a library of transgenic Gal4 enhancer trap lines (Bergeron et al., 2012, 2015). Fish expressing Gal4 in specific neuronal populations were crossed to transgenic UAS:epNTR-RFP, a variant of nitroreductase that converts the prodrug metronidazole into a cell-specific toxin (Horstick et al., 2015; Marquart et al., 2015) (Fig 2A). We screened 31 Gal4 lines and recovered three lines (*y318-Gal4*, *y334-Gal4*, and *y405-Gal4*) where vibration-induced arrest was diminished in ablated larvae (Fig 2B-C, Fig S3A; full 3D expression can be visualized at zbbrowser.com). Ablations did not affect baseline activity in *y318-Gal4* or *y334-Gal4*, whereas spontaneous swimming was reduced in *y405-Gal4* (Fig 2C, S3B). All three lines showed a similar recovery time to controls (Fig S3C), suggesting that the underlying neurons initiate arrest rather than regulating duration. Furthermore, disruption of vibration-induced arrest did not generalize to electric shock-induced freezing (Fig S4) indicating that vibration-induced arrest was independent of a previously described pathway for freezing behavior (Agetsuma et al., 2010; Duboué et al., 2017).

**Figure 2:**
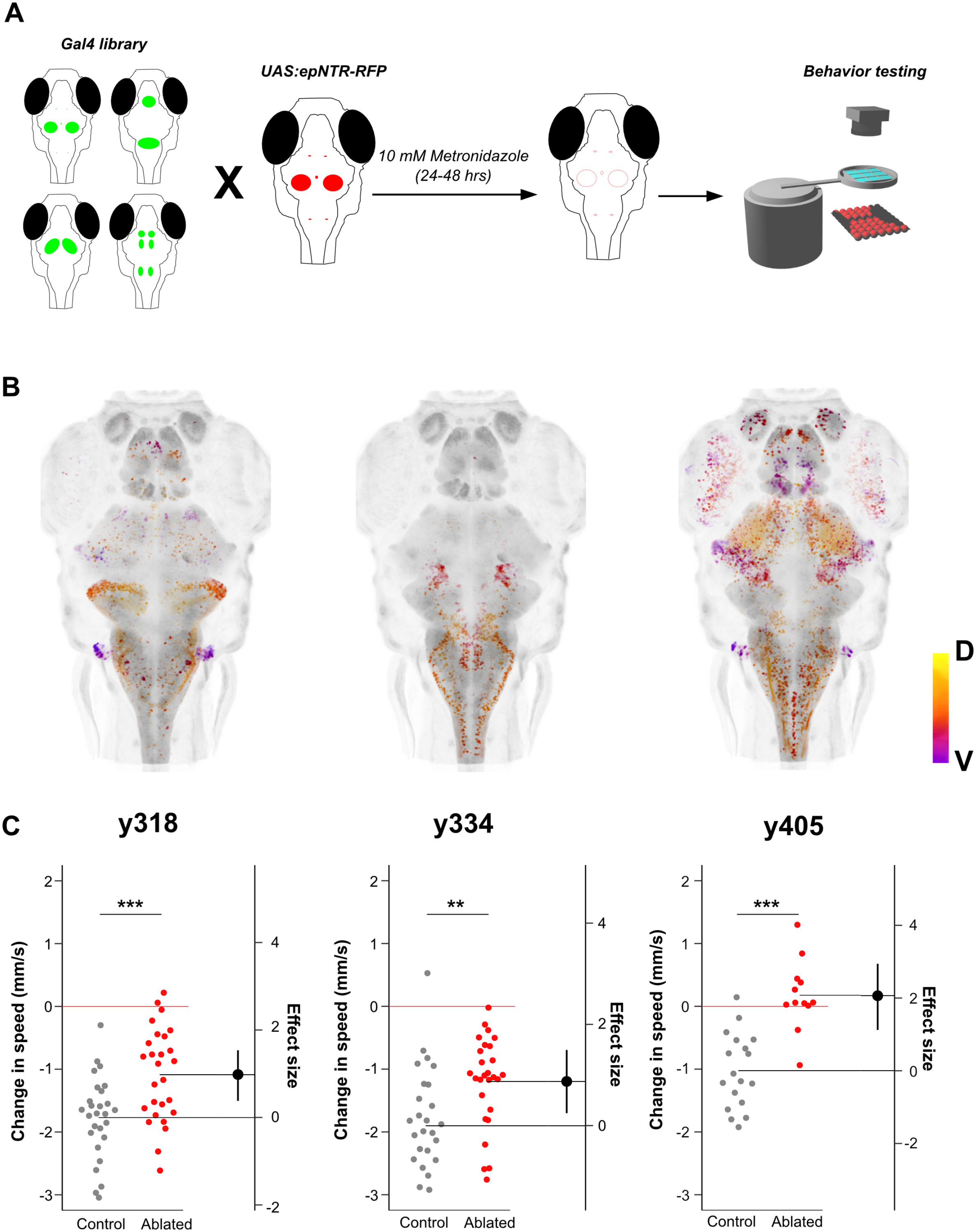
A genetic circuit-breaking screen identifies three lines that contain neurons required for evoking arrest. A Schematic diagram for circuit-breaking screen using 31 enhancer-trap Gal4 lines. Fish were crossed to UAS:epNTR-RFP and treated with 10 mM metronidazole for 24-48 hours to ablate discrete sets of neurons. Fish were tested 24 hours after ablation at 6 dpf for vibration-evoked arrest. B Dorsoventral projection of Gal4 patterns in three lines, *y318-Gal4* (Left), *y334-Gal4* (Middle) and *y405-Gal4* (Right) which showed disruption of arrest following stimulation. Projection is pseudocolor depth coded, D = Dorsal, V = Ventral. All three lines show HuC expression (grey) as counter-label. C Arrest disruption in lines shown in [B]. Arrest defined as the change in speed (mm/s) between baseline and the first epoch following stimulation in ablated (red) and sibling controls (grey). *y318-Gal4* (n = 23 control, 26 ablated), *y334-Gal4* (n = 21 control, 24 ablated), *y405-Gal4* (n = 18 control, 12 ablated). ** p < 0.01, *** p < 0.001, independent samples t-test

### Mechanosensory inputs induce arrest via a cerebellar pathway

Each of the three Gal4 lines labeled neurons in multiple brain regions. To isolate the arrest-related neurons within these lines, we first analyzed *y318-Gal4*, which had prominent expression in the cerebellum accompanied by sparse clusters in other brain regions. We used an intersectional method to selectively label and ablate the cerebellar cluster by crossing *y318-Gal4*;UAS:flox-GFP-epNTR fish to ***y520****-Cre* which has prominent overlapping expression only in the cerebellum (Tabor et al., 2019). Ablation of cerebellar neurons in *y318-Gal4* (Fig. 3A-B) significantly reduced behavioral arrest following vibratory stimuli (Fig 3C) with a similar effect size to complete *y318-Gal4* ablation, indicating that the cerebellar neurons are the relevant subpopulation. *y318-Gal4* cerebellar neurons colocalized with NeuroD-eGFP, a known marker of granule cells (Takeuchi et al., 2015; Volkmann et al., 2008) (Fig 3D) and were largely distinct from the gad1b expressing cerebellar neurons (Fig 3E). These data identify *y318-Gal4* cerebellar granule cells as a component of the arrest-induction circuit.

**Figure 3:**
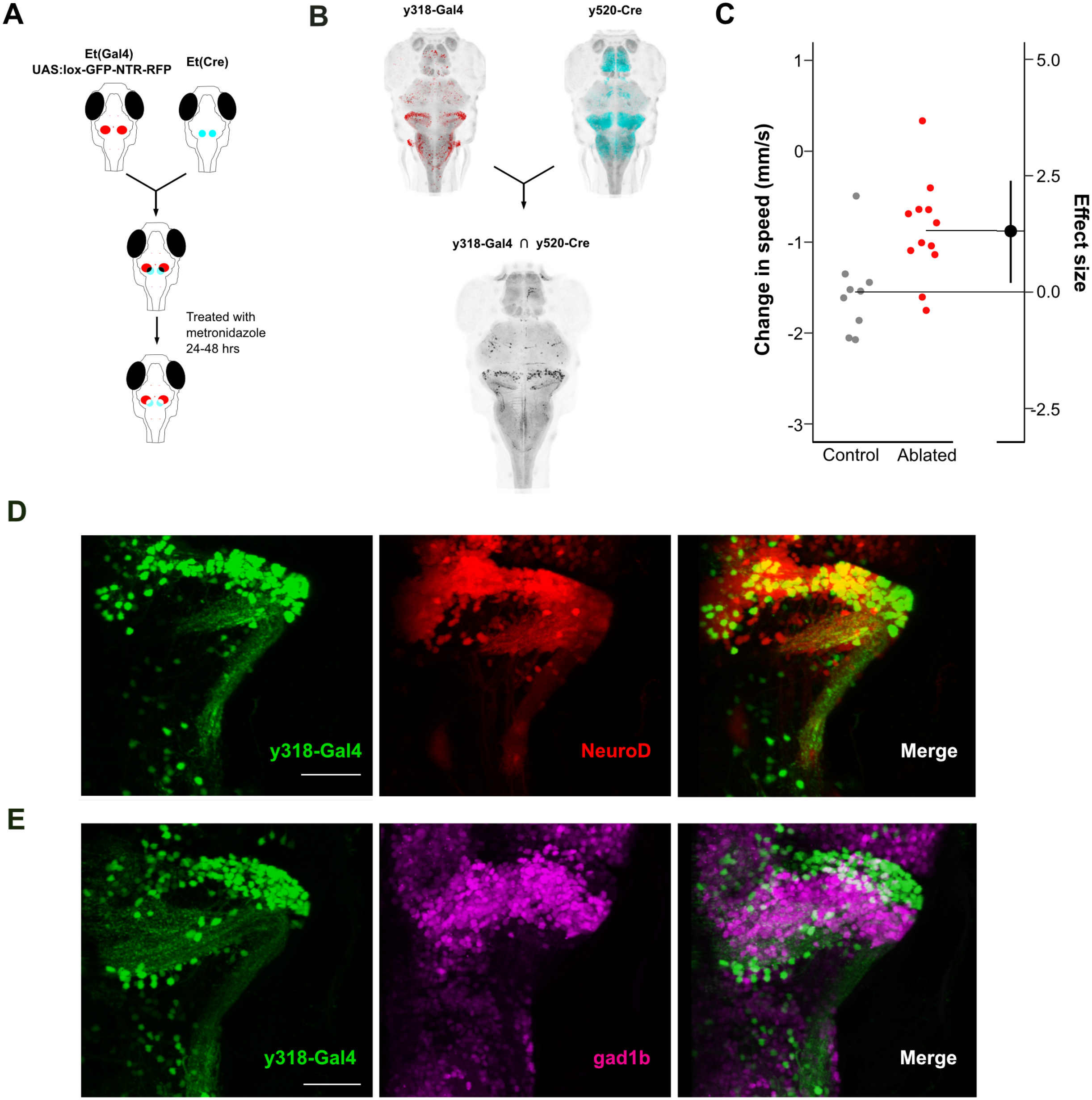
Cerebellar granule cells in *y318-Gal4* are part of the arrest circuit. A Schematic of intersectional ablation approach using Gal4 and Cre to subdivide *y318-Gal4* and spatially restrict NTR expression and ablate Gal4/Cre intersect neurons. B Maximum projections of *y318-Gal4* (red) and *y520-Cre* (cyan) expression patterns. (Bottom) Neurons labeled by NTR in *y318-Gal4*/*y520-Cre* intersect (n = 3 fish). C Ablated *y318-Gal4*/*y520-Cre* intersect neurons (red, n = 12 fish) show reduced locomotor suppression compared to controls (grey, n = 9 fish). Wilcoxon Rank-Sum test p = 0.01. D Maximum projection of *y318-Gal4*;UAS:kaede (green) and NeuroD-eGFP (red) expression shows colocalization of cerebellar *y318-Gal4* neurons and NeuroD (merge). Kaede was photoconverted prior to imaging. Scale bar: 50 μm E Maximum projection of *y318-Gal4*;UAS:kaede (green) and gad1b-RFP (magenta) shows that cerebellar *y318-Gal4* neurons are separate from GABAergic neurons. Scale bar: 50 μm

Involvement of the cerebellum in initiating arrest led us to investigate input and output pathways. Granule cells receive direct input from auditory/vestibular and lateral line ganglia in multiple fish species (Dohaku et al., 2019; Maruska and Tricas, 2009; McCormick et al., 2016; New and Northcutt, 1984). Given that mechanosensory stimuli were used to evoke arrest, we asked whether primary mechanosensory afferents also project to the cerebellum in zebrafish. We labeled afferents by expressing UAS:eGFP-CAAX, a marker for cell membranes, in *y397-Gal4,* which labels the flow sensing anterior and posterior lateral line ganglia, and *y256-Gal4*, which labels part of the auditory/vestibular processing posterior statoacoustic ganglion. As described in previous studies, auditory (Fig 4A-B) and lateral line (Fig 4C-D) afferents both terminated in close proximity to the granule cells of the eminentia granularis (Fig 3E).

**Figure 4:**
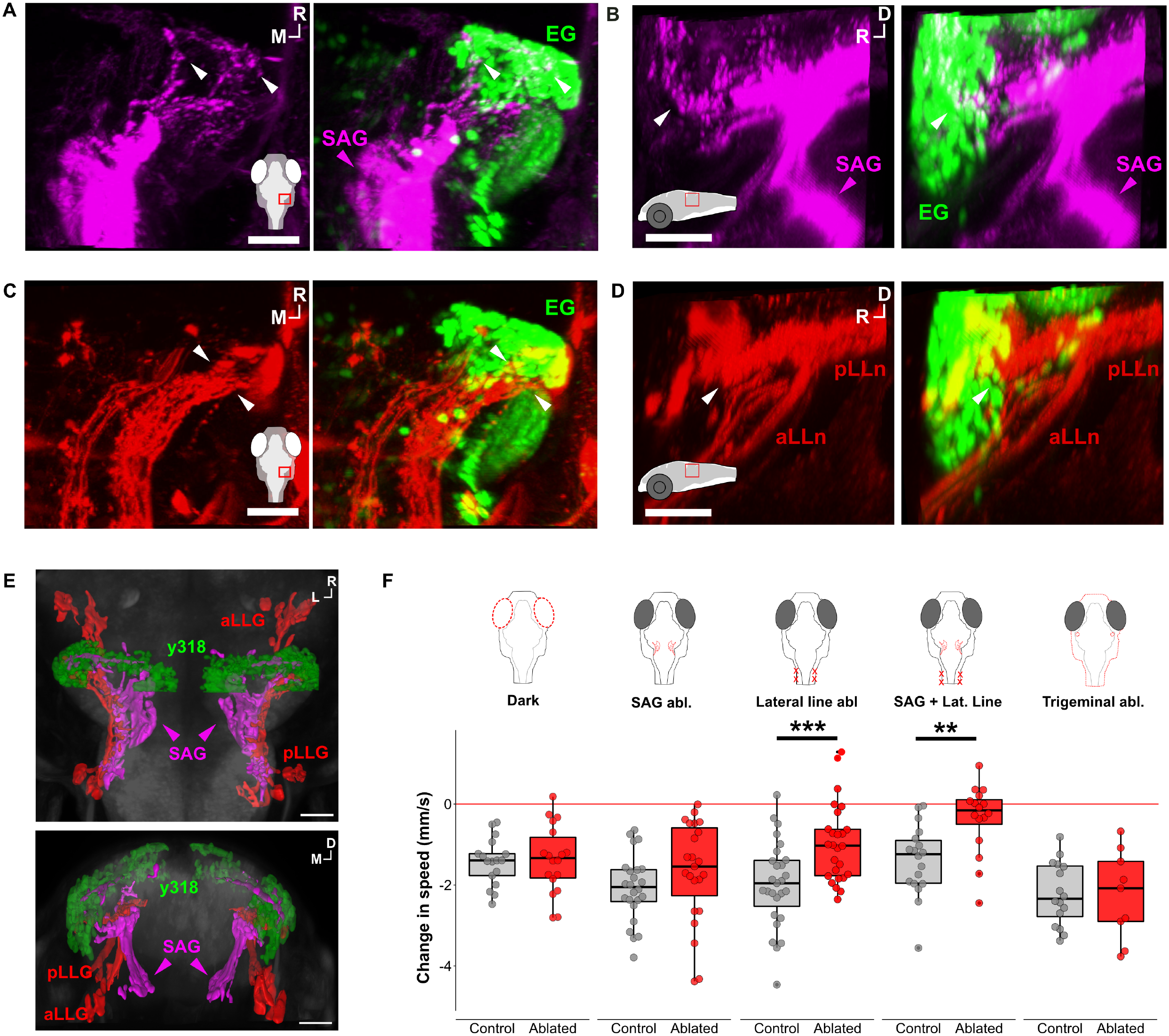
Acoustic and lateral line inputs converge onto lateral cerebellar granule cells to drive arrest. A-B Dorsal and lateral view of *y256-Gal4*;UAS:eGFP-CAAX labeled neurons (magenta) from the posterior statoacoustic ganglion (SAG, magenta arrow) which send projections that terminate in the eminentia granularis (EG) of the cerebellum (white arrows). (Right) Overlay of *y318-Gal4* neurons with SAG neurons. Inset: location of imaged area. Scale bar: 30 μm C-D Dorsal and lateral view of *y397-Gal4*;UAS:eGFP-CAAX labeled neurons (red) from anterior (aLLn) and posterior (pLLn) lateral line which send projections that terminate in the EG (white arrows). (Right) Overlay of *y318-Gal4* neurons with lateral line neurons. Inset: location of imaged area. Scale bar: 30 μm E. Dorsal (Top) and coronal (Bottom) views of 3D reconstruction of SAG neurons from [A] (magenta) and neurons from the anterior lateral line ganglion (aLLG) and posterior lateral line ganglion (pLLG) (red) show that both inputs converge to the EG labeled by *y318-Gal4* neurons (green). D = Dorsal, M = Medial, R = Rostral. Scale bar: 50 μm. F. Effects of removing visual cues (Dark; n = 17 control, n = 18 dark), auditory cues via *y256-Gal4* ablation (SAG Abl, n = 23 control, n = 24 ablated), lateral line cues with 200 μM neomycin treatment (Lateral line abl, n = 27 control, n = 26 ablated), combined SAG and lateral line cues (n = 18 control, n = 16 ablated), and somatosensory cues using *y234-Gal4* ablation (Trigeminal abl, n = 16 control, n = 9 ablated) on arrest following vibratory stimulus. Lateral line ablation suppressed arrest (F_1,52_ = 14.28, p = 0.0004) and combined SAG and lateral line ablation also suppressed arrest (F_1,33_ = 11.98, p = 0.001). *** p < 0.001, ** p < 0.01

Afferent inputs from the lateral line and auditory system into the cerebellum suggested that these sensory modalities are required for evoking arrest. As our arrest-inducing stimulus potentially activated multiple sensory modalities, we tested the role of visual, acoustic/vestibular, lateral line, and somatosensory systems. We disrupted auditory and vestibular inner ear function by ablating posterior statoacoustic ganglion neurons labeled in *y256-Gal4* with UAS:epNTR-RFP and found no effect on arrest (Fig 4F; Fig S5A). We then ablated the flow-sensing lateral line neuromasts using bath application of 250 uM neomycin (Fig S5B). Neomycin ablation reduced arrest by 30% compared to non-treated controls. However, combined ablation of posterior auditory afferents and lateral line neuromasts led to a total loss of arrest, indicating that both flow and acoustic/vestibular information coordinately drive arrest. In contrast, arrest was induced normally when we conducted the experiment in darkness to remove visual cues or disrupted somatosensory inputs. These data indicate that inner ear and lateral line signals evoke behavioral arrest via inputs to eminentia granularis granule cells.

### Glutamatergic prepontine neurons are critical for arrest

Having established that inner ear and lateral line signaling to cerebellar granule cells forms part of the circuit that initiates arrest, we searched for other components of the circuit by examining arrest-related neurons within *y405-Gal4* and *y334-Gal4*. Although these lines express Gal4 in multiple brain regions, a cluster of ventral prepontine neurons (vPPNs) were labeled in both, making them strong candidates (Fig 5A). Targeted multiphoton laser ablation of vPPNs significantly decreased arrest (Fig 5B-C) in a similar magnitude to *y334-Gal4* ablation (Fig 2C).

**Figure 5:**
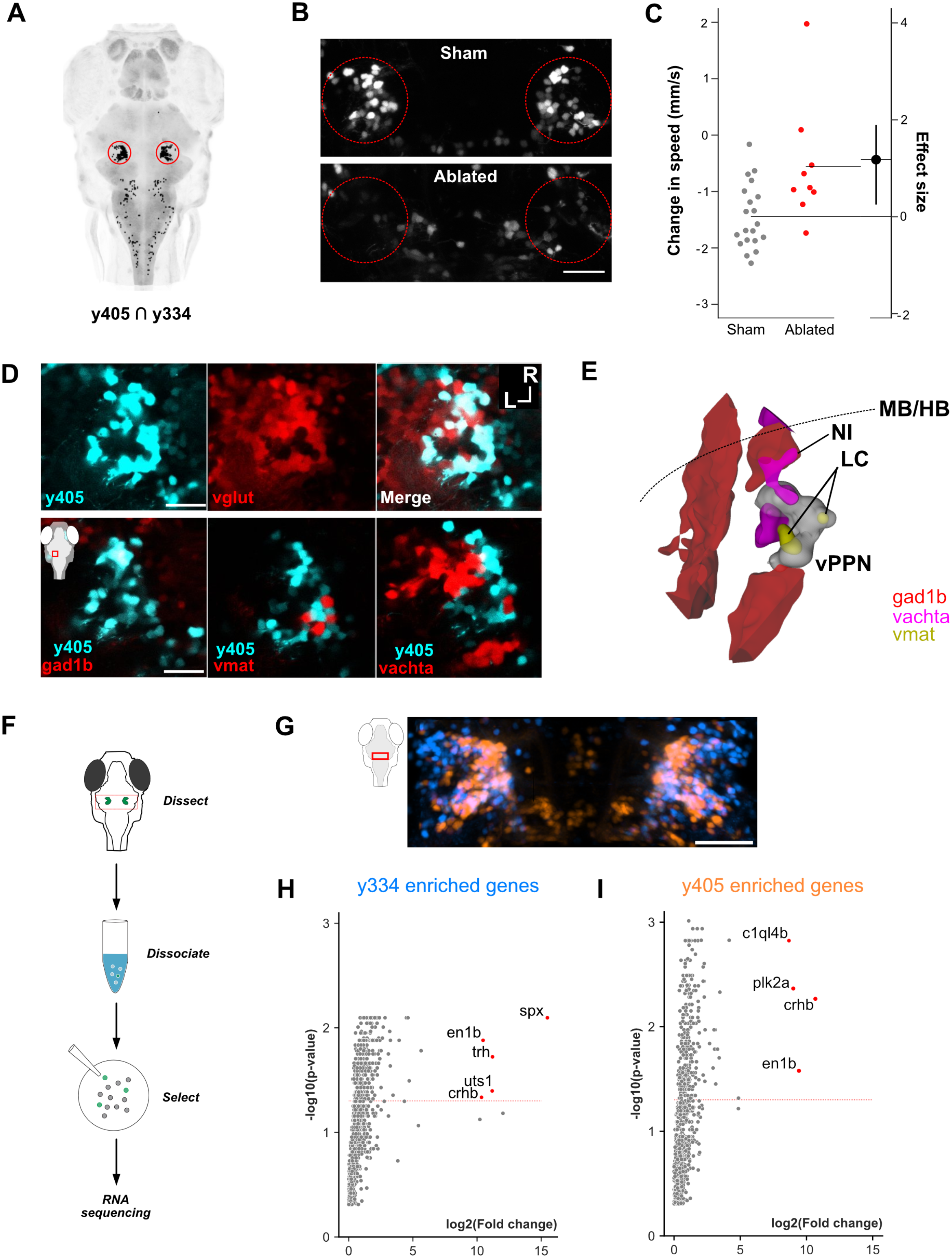
Multiple neuropeptide-expressing glutamatergic vPPNs in *y334-Gal4* and *y405-Gal4* are essential for triggering arrest. A. Computational predicted overlap between *y334-Gal4* and *y405-Gal4* (black pixels) shows a common set of prepontine neurons (outlined by red circles). HuC (grey) used as counter-label. B. Example of multiphoton ablation of *y405-Gal4* vPPNs showing maximum projections of sham ablated vPPNs (Top) and bilaterally ablated vPPNs (Bottom). Circles highlight approximate boundaries of vPPN. Scale bar: 40 μm C. Behavioral arrest in control (grey, n = 20 fish) and *y405-Gal4* ablated (red, n = 10 fish) shown as change in speed following vibrational stimuli. Wilcoxon Rank Sum test, p < 0.001. D. (Top) Overlay of *y405-Gal4*;UAS:kaede (cyan), glutamatergic neurons specified by vglut-GFP (red), and overlay of both. Kaede was photoconverted before imaging. R = Rostral, L = lateral. Scale bar: 25 μm. (Bottom) Overlay of *y405-Gal4* (cyan) with gad1b (Left), vmat2 (Middle), and vachta (Right) neurons. Inset: schematic of imaging window relative to zebrafish brain. Scale bar: 25 μm. E. Dorsal view of 3D reconstruction of *y405-Gal4* vPPNs (white), noradrenergic neurons of the locus coeruleus (LC) labeled by vmat2-GFP (yellow), cholinergic neurons of the nucleus of the isthmus (NI), and GABAergic neurons (red). Imaging was conducted on individual lines and registered to a common reference before analysis. Dotted line shows midbrain-hindbrain boundary (MB/HB). F. Schematic of selected neuron RNA sequencing used to identify neurotransmitter identity of vPPNs. vPPNs from *y334-Gal4*/*y405-Gal4* are dissected, dissociated, and plated onto a petri dish. Following visual selection for kaede-expressing neurons, samples undergo library preparation and RNA sequencing. G. Maximum projection of *y334-Gal4* (blue) and *y405-Gal4* (orange) expression in vPPNs. Schematic shows approximate imaging window in the fish brain. Scale bar: 40 μm H. Volcano plots of RNA-seq data showing genes enriched in *y334-Gal4* [H] and *y405-Gal4* [I] relative to a pan-neuronal reference. Labeled genes (red) show log_2_ fold changes > 8 and FDR-corrected p-values < 0.05. Dashed red line shows threshold of Benjamini-Hochberg corrected 5% false discovery rate.

vPPNs are located in a rostral part of the hindbrain that comprises multiple neuronal cell groups (Kinkhabwala et al., 2011; Watson et al., 2019), including glutamatergic, GABAergic, and cholinergic neurons and the noradrenergic locus coeruleus. vPPN neurons labeled by *y334-Gal4* and *y405-Gal4* showed considerable overlap, but *y334-Gal4* had greater expression in the rostral and lateral vPPN (Fig 5G). To further characterize vPPNs, we performed colocalization imaging experiments with transgenic lines that label major neurotransmitters (Fig 5D). The majority of vPPNs (97%, 288/296 cells from n = 5 fish) colocalized with vglut2a:GFP indicating that these neurons are primarily glutamatergic. vPPNs did not colocalize with GABAergic neurons labeled by gad1b:dsRed and were adjacent to, but did not overlap with, cholinergic (vachta:GFP) neurons. Intriguingly, cells expressing vmat2:GFP, which labels the locus coeruleus, were interspersed with caudal vPPNs but we did not find vPPNs that co-expressed GFP (n=108 neurons, Fig 5D-E). To better understand vPPN identity, we analyzed transcriptomes of prepontine neurons from *y334-Gal4* and *y405-Gal4* using RNA sequencing (Fig 5F, H-I). Confirming that we correctly isolated the prepontine neurons from these lines, both *y334-Gal4* and *y405-Gal4* transcriptomes showed enrichment for engrailed 1b (eng1b), which is expressed at the midbrain-hindbrain boundary (Ekker et al., 1992). Among the most highly enriched transcripts in vPPNs were several stress-associated neuropeptides, including corticotropin releasing hormone b (crhb), thyrotophin releasing hormone (trh), and urotensin 1 (uts1) (Fig 5H-I). Together, these data define a novel area for defensive responses in the prepontine tegmentum, consisting of glutamatergic cells that coexpress multiple neuropeptides, that is bounded caudally by the locus coeruleus and laterally by cholinergic cells of the nucleus isthmi (Henriques et al., 2019)(Fig 5E).

### Cerebellar Purkinje neurons project to prepontine arrest neurons

vPPNs were positioned in the tegmentum directly below the cerebellum raising the possibility of a direct connection as part of the arrest circuit. We therefore examined efferent projections of Purkinje cells in *aldoca-Gal4;*UAS:eGFP-CAAX transgenic larvae. Along with the previously reported caudal projection to the statoacoustic ganglion (Bae et al., 2009; Matsui et al., 2014) (Fig. 6A, yellow arrows), we noticed a small rostroventral projection from lateral *aldoca-Gal4* neurons. These Purkinje cell projections terminated within the caudal vPPN (Fig 6A, white arrows) and suggested that Purkinje cells are presynaptic partners of vPPNs. Because Purkinje cells are GABAergic, we asked whether vPPNs receive inhibitory inputs. Labelling the post-synaptic inhibitory synapses formed by vPPN neurons using ***y334-*** *Gal4*, UAS:gephyrin-FingR-mCherry transgenic larvae (Son et al., 2016) revealed that a subset of vPPNs, in a similar caudal region to the area receiving Purkinje neurons projections, showed mCherry fluorescence (Fig 6B, white arrows), supporting the idea that these neurons receive inhibitory input. We therefore pharmacologically manipulated GABAergic signaling. Treatment with either GABA or ethanol, an indirect GABA-A receptor agonist, reduced vibration-induced behavioral arrest (Fig. 6C-D). After ethanol washout, larvae recovered fully within 20 min. Because bath applied treatments likely affected GABA signaling throughout the brain, we tested whether blocking GABA_A_ receptor signaling specifically in vPPNs suppressed arrest. To do so, we generated a UAS:ICL-GFP construct to express the intracellular loop (ICL) from the GABA_A_ receptor γ2 subunit in vPPNs (Fig 6E). Expression of this peptide in Xenopus reduces GABA_A_ currents resulting in decreased GABA-mediated behaviors (Shen et al., 2009, 2011). *y334-Gal4* fish expressing UAS:ICL-GFP showed reduced behavioral arrest (Fig 6F), whereas controls that expressed a mutant form of the ICL that does not block GABAergic signaling (Shen et al., 2009) responded normally (Fig 6F). Together, these data suggest that GABAergic inhibition of vPPN neurons, potentially by direct input from Purkinje cells, plays a central role in evoking behavioral arrest in response to an overwhelming threat.

**Figure 6:**
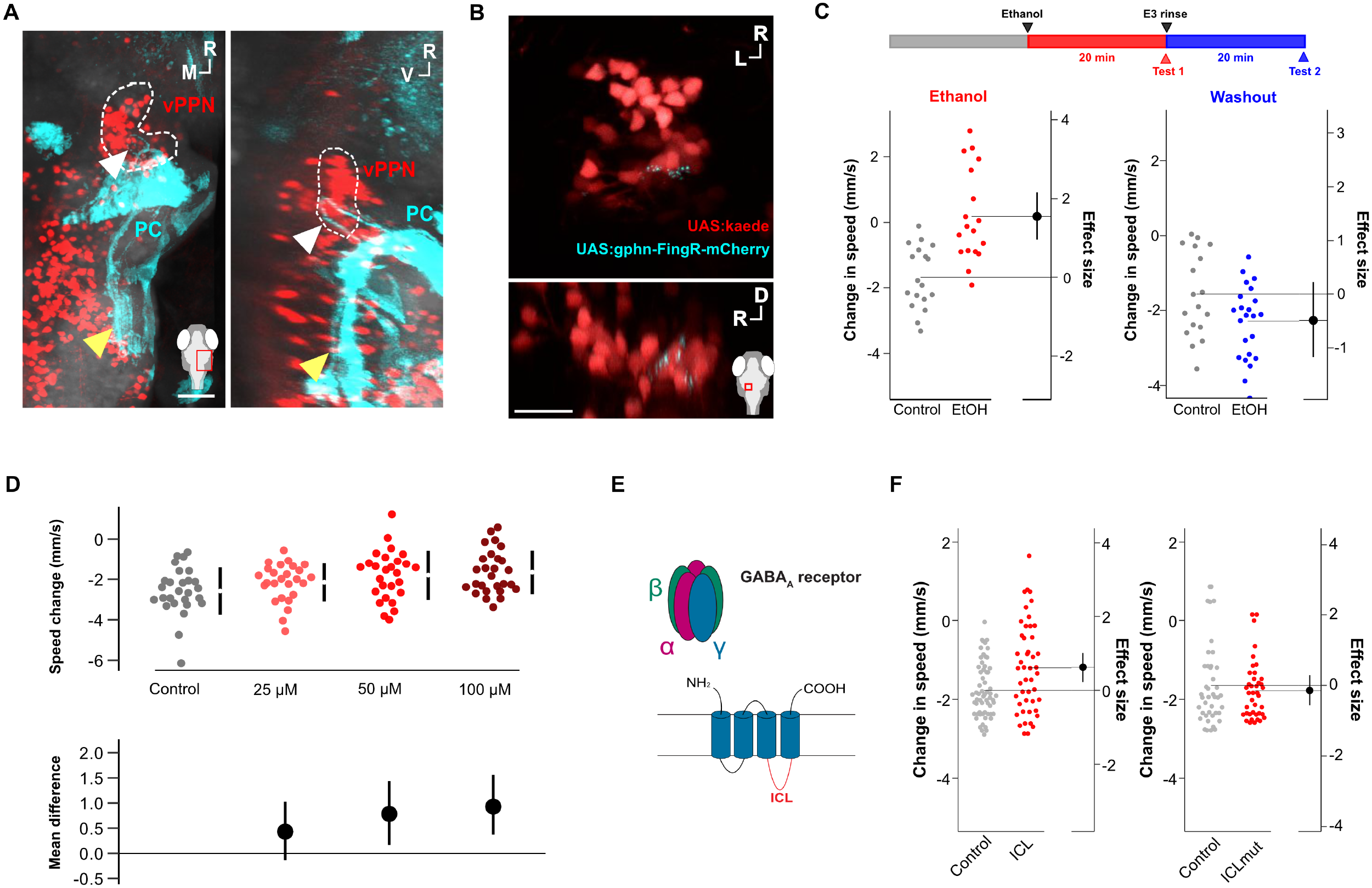
GABAergic signaling from Purkinje cells to vPPNs regulates arrest onset. A Dorsal (Left) and lateral (Right) views of maximum projections of Purkinje cells (PC) labeled by *aldoca-Gal4*;UAS:eGFP-CAAX (cyan) and *y334-Gal4* vPPNs (outlined with dashed white line) following co-registration. White arrows show projections from PCs toward vPPNs and yellow arrows show hindbrain projections. R = Rostral, M = Medial, V = Ventral. Inset shows imaging window. Scale bar: 50 μm B Dorsal (Left) and lateral (Right) views of maximum projections of *y405-Gal4* vPPNs colabeled with UAS:kaede (red) and UAS:gphn-FingR-mCherry (cyan). R = Rostral, L = Lateral, D = Dorsal. Inset shows imaging window. Scale bar: 20 μm C (Top) Schematic diagram of experimental protocol for ethanol exposure experiment showing timepoints for testing ethanol exposed fish (red) and washout experiment (blue). (Bottom left) Behavioral arrest in control (grey, n = 18 fish) and 300 mM ethanol exposed (red, n = 18 fish) shown as change in speed following vibrational stimuli. Two sample t-test, p < 0.001. (Bottom right) Behavioral arrest in control (grey, n = 18 fish) and ethanol washout (blue, n = 18 fish) shown as change in speed following vibrational stimuli. Two sample t-test, p = 0.72. D Change in behavioral arrest after bath application of GABA at concentrations of 25 µM (pink, n = 27 fish), 50 µM (red, n = 27), and 100 µM (dark red, n = 27) compared to control (grey, n = 27). Top row shows raw change in speed and bottom row shows 95% CI of mean differences relative to control. ANOVA, p = 0.01. E Schematic representation of the GABA_A_ receptor complex with the intracellular loop of the γ subunit (ICL). UAS:ICL-GFP expression suppresses GABA_A_ signaling and the mutant form of ICL (UAS:ICLmut-GFP) does not affect GABA_A_ signaling. F Changes in vibration-evoked arrest in *y334-Gal4* fish (Left) expressing UAS:ICL-GFP (grey = injected controls, n = 61; red = ICL-expressing, n = 47; two-sample t-test, p = 0.007) and (Right) expressing UAS:ICLmut-GFP (grey = injected controls, n = 42; red = ICLmut-expressing, n = 40; two-sample t-test, p = 0.91).

### Prepontine arrest neurons increase activity following persistent flow stimuli

To test whether vPPNs respond to threat, we measured changes in phosphorylated ERK (pERK), a marker for neural activity, using immunohistochemical labeling after exposure to the intense vibratory stimulus (Randlett et al., 2015). After exposure, larvae were fixed, labeled, imaged, and co-registered. Stimulation led to increased pERK/tERK ratios in the posterior tuberculum, midbrain tegmentum and prepontine regions including in the area occupied by vPPNs (Fig 7A-B). However, because changes in pERK lag neuronal activity by around 2 min (Dai et al., 2002), it was possible that the pERK signal reflected peri-stimulus activation preceding behavioral arrest. We therefore examined vPPN activity to intense flow stimuli at higher time resolution by expressing a nuclear localized UAS:GCaMP6s in *y334-Gal4* and measuring changes in fluorescence following an intense pulsed flow stimulus in a head-fixed preparation (Fig 7C-E). A subset of vPPNs (n = 41/420 neurons from 8 fish) showed a strong increase in fluorescence after several seconds of flow stimulation (Fig 7D-E). Cells with increased GCaMP6s fluorescence were highly stereotyped and robust in their responses to successive stimuli (Fig S7). The increase in GCaMP6s fluorescence was delayed and GCaMP6s fluorescence peaked in stimulus-responsive cells at the cessation of the stimulus. This delayed increase cannot be accounted for by nuclear-localized GCaMP6s kinetics, which peaks ∼1.1 s after activation (Förster et al., 2017) and therefore suggests that vPPN activity builds during exposure to an overwhelming threat.

**Figure 7:**
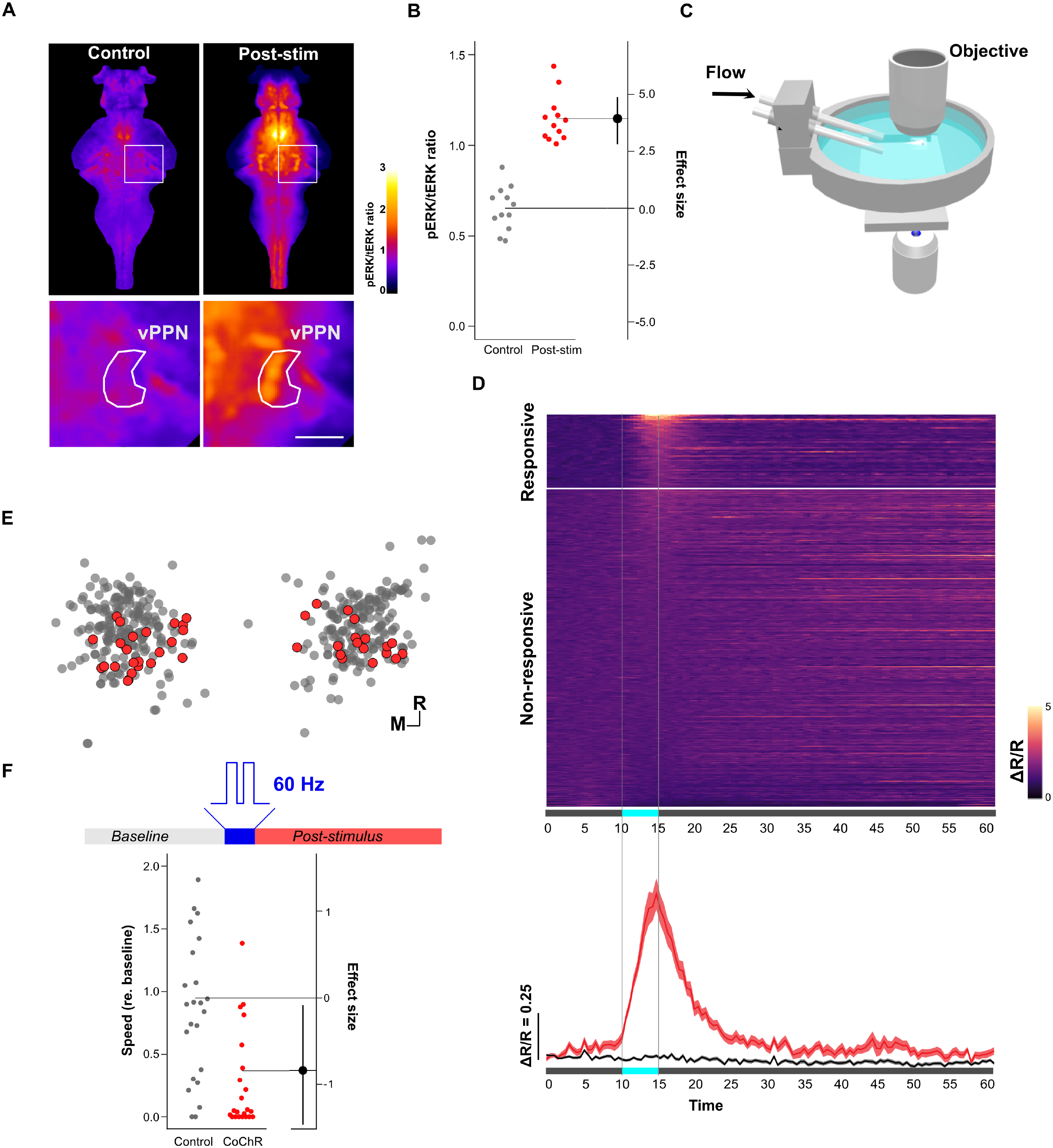
vPPN activation is associated with inescapable flow stimulation and drives arrest. A (Top) Single plane dorsal view of registered, averaged pERK/tERK ratios in control (n = 12 fish) (Left) and vibratory-stimulus exposed fish (n = 12) (Right). Color shows p/tERK ratio (AU). (Bottom) Magnified region around right vPPN showing greater activation in post-stimulus condition compared to control. Scale bar: 30 μm B pERK/tERK ratios in vPPN masked regions for control (grey, n = 12 fish) and vibration stimulus exposed (red, n = 12 fish) conditions. ANOVA F_1,22_ = 92.98, p < 0.001. C Schematic of apparatus used to deliver water flow stimuli to head-fixed zebrafish for calcium imaging. D (Top) Calcium traces for flow-responsive (n = 41 cells from 14 fish) and non-responsive (n = 379 cells) neurons showing GCaMP6s fluorescence normalized to nuclear-localized RFP (ΔR/R). Vertical dashed line shows flow stimulation period. (Bottom) Average (± SEM) normalized GCaMP6s fluorescence for responsive (red) and non-responsive (black) neurons. E Position of flow-responsive (red) and non-responsive (grey) neurons within the vPPN. R = Rostral, M = Medial. F (Top) Schematic diagram of photostimulation protocol with 60 Hz 2 W/cm^2^ stimulus. (Bottom) Behavioral arrest represented as speed (relative to baseline) in injected controls (grey, n = 23 fish) and *y334-Gal4*;UAS:CoChR (red, n = 22 fish) following photostimulation. ANOVA F_1,45_ = 8.196, p = 0.006.

If vPPNs were activated during arrest-inducing stimuli, we reasoned that activating vPPNs would evoke arrest in the absence of vibrational stimuli. We expressed the channelrhodopsin variant UAS:CoChR, which has been shown to produce robust activation in zebrafish neurons (Antinucci et al., 2020). Photoactivation in free swimming *y334-Gal4*;UAS:CoChR fish (Fig 7F) resulted in significant locomotor suppression after photoactivation, whereas sibling controls lacking UAS:CoChR showed no change in motor activity (Fig S8). Consistent with vPPN ablation eliminating arrest, vPPN activation simulated exposure to an intense threat and induced arrest.

### Prepontine neurons contact premotor reticulospinal neurons

The critical role of vPPNs in evoking arrest led us to map efferent targets of vPPNs. We labeled individual *y334-Gal4* neurons using an intersectional approach combining UAS:blo-GFP-blo-lynTagRFP and a heat shock-inducible B3 recombinase (Tabor et al., 2018), taking advantage of the inefficiency of B3 recombinase in zebrafish to facilitate sparse labeling (Fig 8A). After imaging, registering, and reconstructing these neurons (13 neurons from 5 fish), we identified three classes of vPPNs, two of which projected ventrally to a dense neuropil region (Fig 8B). The first class (represented by n = 7 neurons) projected ventrally to an ipsilateral neuropil region between the hypothalamus and the tegmentum. We noted that these projections travelled, within the margin of registration error, adjacent to the cell body of RoL1 neurons, a cluster of reticulospinal neurons that project to the spinal cord (Gahtan and O‘Malley, 2003) that drive forward locomotor bouts and stimulus-evoked turns (Lovett-Barron et al., 2020; Orger et al., 2008) (Fig 8B). A second class of vPPNs (n = 5 neurons) projected ventrally to the ipsilateral RoL1s and also crossed the midline to terminate near the contralateral RoL1 cluster. Contralateral projections were ordered rostrocaudally with rostral vPPN axons terminating in the rostral contralateral RoL1. The third class (n = 2 neurons) projected anteriorly to the hypothalamus and torus semicircularis. To test whether vPPN termini contacted RoL1 neurons, we expressed UAS:synaptophysin-RFP in *y334-Gal4* and backfilled RoL1s using fluorescently labeled dextran injected into the spinal cord. Fluorescent puncta from vPPN presynapses colocalized with RoL1cell bodies (Fig 8C-D). We also observed numerous presynaptic puncta in the neuropil region surrounding RoL1 neurons.

**Figure 8:**
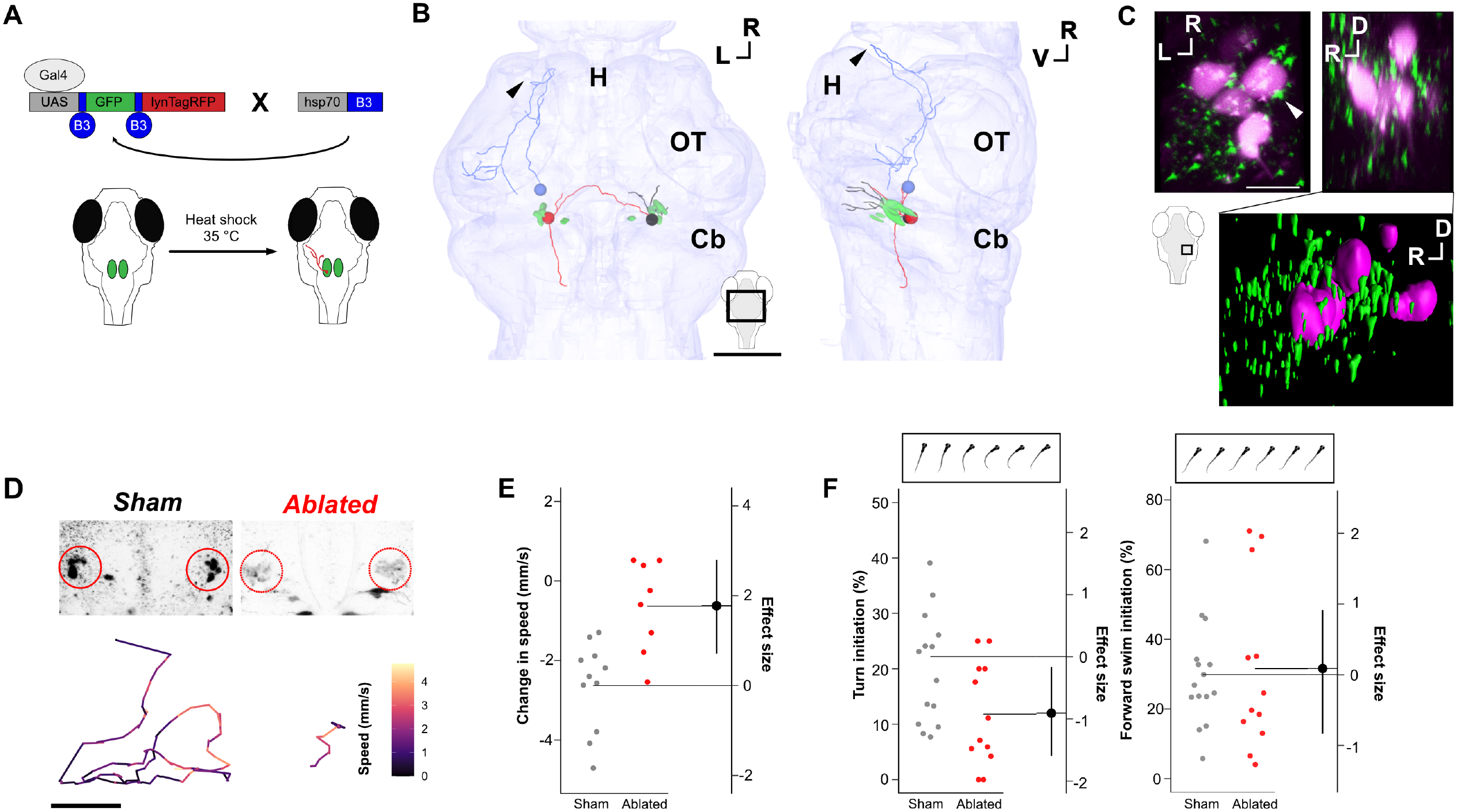
vPPN neurons project bilaterally to RoL1 reticulospinal neurons and disrupt their activity to evoke arrest. A Diagram of Gal4/UAS and B3 recombinase intersectional approach used for single neuron tracing. Heat shock activates B3 recombinase and facilitates sparse labeling of single neurons with membrane-tagged RFP (lynTagRFP) B Dorsal (Left) and lateral (Right) views of maximum projection of single neuron traces from *y334-Gal4* vPPNs in a 3D model of zebrafish brain with a model of RoL1 reticulospinal neurons (green). Spheres denote cell body location. Representative examples of vPPNs projecting to ipsilateral (black), contralateral (red) and hypothalamic (blue) targets are shown (Arrow shows terminal projection in rostral hypothalamus. All neurons can be viewed in Figure S6. Inset shows viewing window. Scale bar: 50 um. R = Rostral, L = Lateral, V = Ventral. H = Hypothalamus, OT = Optic Tectum, Cb = Cerebellum C (Top) Maximum projection of *y334-Gal4*;UAS:synaptophysin-RFP (green) and backfilled RoL1 (magenta) colocalization in Dorsal (Left) and Lateral (Right) views. White arrow shows example of apposition of *y334-Gal4* synapses with RoL1 cell body. Inset: imaging window. Scale bar: 10 um. (Bottom) 3D rendering of lateral view showing synapses relative to RoL1 and surrounding neuropil. R = Rostral, L = Lateral, D = Dorsal D (Top) Representative examples of backfilled sham ablated (Left) and multiphoton ablated (Right) RoL1 reticulospinal neurons. (Bottom) Representative baseline movement traces for sham ablated (Left) and RoL1 ablated (Right) fish. Color represents speed within a 5 sec epoch. Scale bar: 1 cm. E Behavioral arrest measured as change in speed (mm/s) for sham ablated (grey, n = 11 fish) and bilateral RoL1 ablated (red, n = 8 fish). ANOVA F_1,17_ = 14.63, p = 0.001. F Turn initiation (Left) and forward swim initiation (Right) measured from high-speed video recordings during baseline locomotion in sham ablated (grey, n = 14) and RoL1 ablated (red, n = 12). Turn initiation is specifically suppressed in RoL1 ablated fish. ANOVA F_1,25_ = 5.49, p = 0.03.

To test whether RoL1s were part of the arrest pathway, we ablated them using a multiphoton laser before testing for changes in locomotor behavior and vibration-evoked arrest. We confirmed previous reports that RoL1 ablation reduced baseline locomotor behavior (Lovett-Barron et al., 2020; Orger et al., 2008) (Fig 8E, S6A). RoL1 ablated fish also showed a loss of behavioral arrest following vibratory stimuli (Fig 8F). Importantly, analysis of swim path trajectories showed that RoL1 ablation not only suppressed overall displacement, but also resulted in straight-line swim trajectories observed during arrest (Fig 8E). Kinematic analysis using high speed recordings showed that RoL1 ablation led to a specific reduction in turn bouts, with forward swims relatively spared (Fig 7F, S6B). These results support a model in which RoL1 neurons promote baseline locomotion that is suppressed via vPPNs in response to severe threat.

## Discussion

Understanding how animals perceive threatening stimuli and rapidly initiate defensive responses is fundamental for understanding cognitive states such as fear (LeDoux, 2000). Our data reveal that larval zebrafish, when presented with an intense and inescapable threat stimulus, respond with behavioral arrest with characteristic features of tonic immobility. As observed in mammals, reptiles, and birds, tonic immobility in zebrafish manifests as an innate response defined by locomotor suppression, lack of responsiveness to external stimuli, and delays in the righting reflex. The magnitude and persistence of this state is intensity dependent as described in chick (Gallup, 1977) and lizards (Edson and Gallup, 1972).We note that tonic immobility has other behavioral and physiological correlates that we did not measure, including decreased heart rate and increased respiration, tremor, and changes in correlated neural activity (Kozlowska et al., 2015; Yoshida, 2021). Further investigation will elucidate whether zebrafish exhibit these physiological changes and may reveal further core conserved elements of this behavior.

Based on our data, we propose a sensory afferent-cerebellar-prepontine circuit in which vPPNs are crucial for initiating arrest following turbulent flow stimuli (Fig 9). In this model, at baseline, vPPNs show low levels of activity allowing RoL1s to propel normal locomotor activity. Transient threats may evoke escape responses, but in the face of persistent danger, combined flow and auditory/vestibular inputs to granule cells stimulate cerebellar activity and increase GABAergic Purkinje cell output. Following prolonged stimulation, loss of GABAergic signaling elicits post-inhibitory rebound firing in vPPNs which in turn suppresses RoL1 firing and therefore disrupts swimming. This model describes a central pathway by which inescapable threatening stimuli induce behavioral arrest in zebrafish and may serve as a model for conserved pathways that mediate tonic immobility in other species.

**Figure 9:**
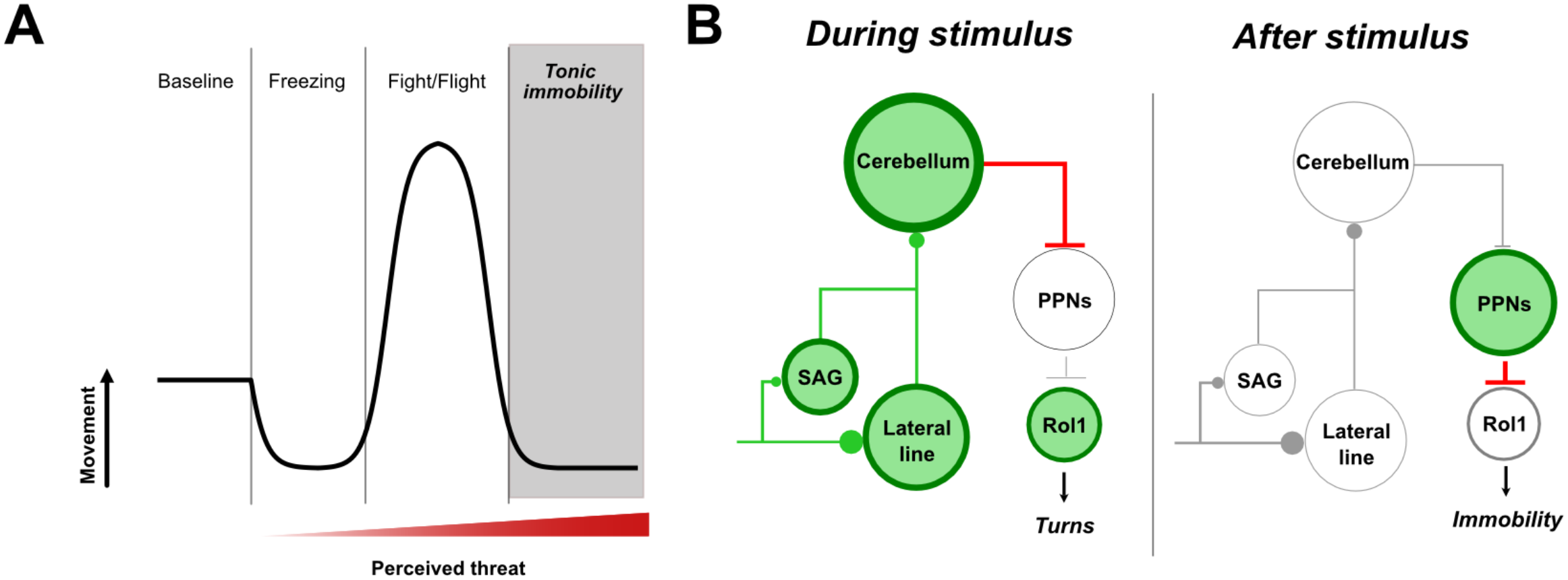
Circuit model for tonic immobility. A Representation of the defense cascade showing changes in activity as a function of perceived threat. The terminal behavior in the defense cascade is tonic immobility, which presents as behavioral arrest (reduced movement compared to baseline) B Repeated, inescapable stimuli activate peripheral auditory (SAG) and lateral line ganglia which then excite the granule cells of the cerebellum (green lines). During stimulus presentation, the cerebellum sends inhibitory projections to the prepontine neurons (PPNs). Following sustained vibratory stimulation, cerebellar inhibition ceases and leads to activation of the PPNs. Bilateral activation of the PPNs disrupts RoL1 neuron activity, leading to suppressed turning behavior and immobility.

In our working model, persistent Purkinje cell inhibition elicits post-inhibitory vPPN activation. GABA facilitation and *y334-Gal4* specific GABA_A_ receptor blockade both disrupted vibration-evoked behavioral arrest (Fig 6C-E). Although these results appear contradictory, we propose that blocking initial GABAergic signaling suppresses inhibition whereas pharmacological GABA facilitation may prevent the sudden offset of inhibition. Both experiments would thus disrupt post-inhibitory firing in vPPNs and lead to the same behavioral outcome. Post-inhibitory rebound firing has been demonstrated in glutamatergic targets of Purkinje cells (Aizenman and Linden, 1999; Witter et al., 2013; Zheng and Raman, 2009), and in thalamocortical neurons (Sohal et al., 2006), suprachiasmatic nucleus of the hypothalamus (Tremere et al., 2008), and amygdala (Ryan et al., 2012). In these circuits, post-inhibitory rebound facilitates synchronization of multiple neuron populations (Ryan et al., 2012; Sohal et al., 2006) similar to the bilateral activation of vPPNs neurons in our experiments (Fig 7D-E). Our model is limited by our incomplete understanding of direct synaptic and functional connectivity between Purkinje cells and vPPNs. Nevertheless, we propose that synchronized post-inhibitory activation after prolonged cerebellar inhibition is a plausible mechanism for arrest and invite future experiments to explicitly test this hypothesis.

How does bilateral activation of glutamatergic vPPNs result in locomotor inhibition? vPPNs project bilaterally to RoL1 reticulospinal neurons (Fig 8B-D) and bilateral RoL1 ablation leads to a decrease in activity similar to vPPN activation (Fig 8E-G). vPPNs show strong expression of several stress-associated neuropeptides including crhb, trh and uts1. Co-release of these neuropeptides with glutamate may suppress or disrupt RoL1 function, leading to behavioral arrest. Alternatively, vPPNs may modify other inputs to RoL1 neurons – RoL1s are located in a dense neuropil that receives projections from many brain regions (Barrios et al., 2020; Lovett-Barron et al., 2020). vPPNs make synaptic contacts within this neuropil (Fig 5C) and may therefore influence other convergent pathways that influence turning behavior.

Our data defines a new region within the prepontine tegmentum that mediates responses to persistent threat. Neurons in the vPPN form a characteristic crescent shape in both *y334-Gal4* and *y405-Gal4* transgenics, but these lines have slightly different domains of expression in this region, potentially explaining why *y405-Gal4* ablation had a greater effect on behavioral arrest initiation than *y334-Gal4* ablation (Fig 2C). The difference and is also consistent with our observation that the caudal vPPN, which contains more *y405-Gal4* neurons, has greater activation following pulsed flow stimuli (Fig 8F). RNA sequencing from *y334-Gal4* and *y405-Gal4* show that vPPNs in both lines are enriched in crhb, the teleost homolog of corticotropin releasing hormone (CRH), which is consistent with *in situ* hybridization studies (Alderman and Bernier, 2009; Chandrasekar et al., 2007). CRH is a prominent regulator of the hypothalamus-pituitary-adrenal axis and CRH receptor activation in rodent amygdala and periaqueductal grey facilitates defensive behaviors, including tonic immobility (Sherman and Kalin, 1988; Spinieli and Leite-Panissi, 2018). However, it is possible that crhb may only identify these neurons and that crhb function may be dispensable for tonic immobility.

vPPNs are bounded by the locus ceruleus and nucleus isthmi (the teleost homolog of the parabigeminal nucleus) (Fig 5D). In mammals a similar neuroanatomical region comprises the parabrachial complex, a structure in the prepontine region lateral to the locus ceruleus that is dissected by the superior cerebellar peduncles (Fulwiler and Saper, 1984; Palmiter, 2018). Intriguingly, the parabrachial nucleus has been implicated in both tonic immobility (Klemm, 2001; Menescal-de-Oliveira and Hoffmann, 1993) and freezing in mouse (Bowen et al., 2020; Han et al., 2015). In mammals, the parabrachial nucleus integrates aversive stimuli and serves as a general alarm system for threats (Barik et al., 2018; Chiang et al., 2019; Palmiter, 2018). Like vPPNs, most parabrachial neurons are glutamatergic and are enriched in CRH expression (Palmiter, 2018). Thus, based on similarity of position, function, and gene expression, we provisionally propose that vPPNs are homologous to a subnucleus of the mammalian parabrachial complex. Future experiments examining molecular markers of the parabrachial nucleus and functional connectivity may provide more direct evidence for homology. If so, our data would indicate that the parabrachial complex has a deep evolutionary history in promoting defensive responses.

Our results identified the cerebellum as a key structure in vibration-induced arrest. Indeed, cerebellar inactivation suppresses freezing and tonic immobility in mammals (Koutsikou et al., 2014; McBride and Klemm, 1969; Supple et al., 1988) and *cerebelless* mouse mutants, which lack GABAergic neurons in the cerebellar nuclei, fail to show tonic immobility when restrained by neck pinch (Esposito et al., 2013). Moreover, the cerebellum is known to encode painful stimuli and modulate defensive responses in rats (Saab and Willis, 2003), suggesting a broader role for the cerebellum in defensive behaviors. Similarly, teleost Purkinje cells show increased tonic firing following a sudden change in the sensory environment (Hsieh et al., 2014). We found that behavioral arrest required coordinated signals from the auditory and lateral line systems. Granule cells of the eminentia granularis receive input from both pathways and are therefore a likely point of sensory integration (Ishikawa et al., 2015; Sawtell, 2010) (Fig 4E). The partial disruption of immobility in *y318-Gal4* granule cell ablation compared to total loss of immobility in combined lateral line and auditory/vestibular ablations may indicate that *y318-Gal4* labels only a subset of granule cells, or that a second pathway processes sensory threats. Understanding how lateral line and auditory/vestibular information converges in granule cells to evoke arrest remains an open question.

Defensive behaviors associated with fear are instances of ‘emotion primitives’, a conceptual framework that defines internal states with behavioral characteristics that scale with intensity, valence, and persistence (Anderson and Adolphs, 2014). This study shows that tonic immobility in larval zebrafish meets these criteria and outlines the underlying circuit. Our data disclose that vPPNs play a central role in behavioral state control by activating the terminal behavior in the defense cascade. Understanding how the vPPNs over-ride expression of other defensive behaviors will help us better understand how animals assess risk and match defensive responses to perceived threat.

## Supporting information

**Figure S1.**
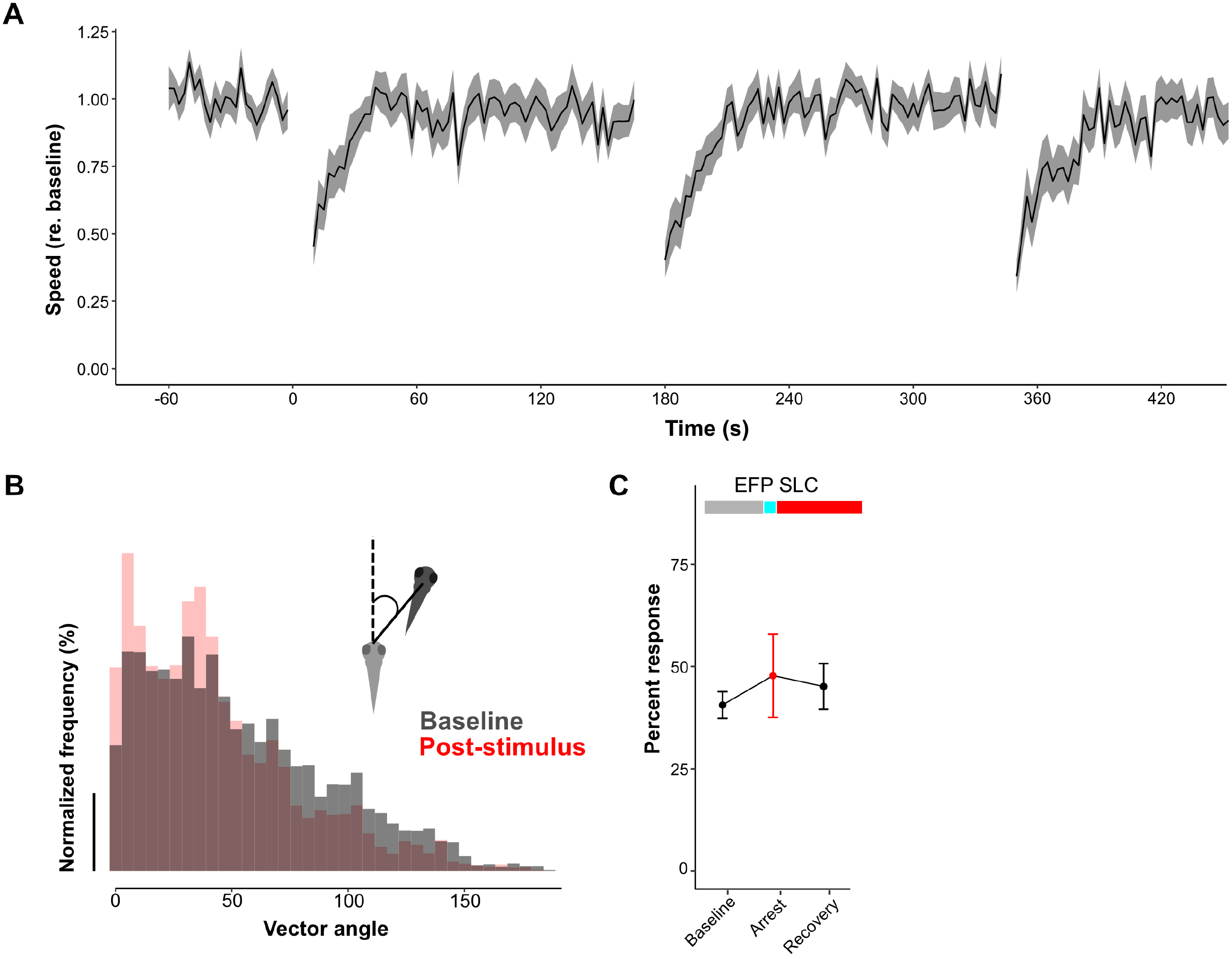
A Averaged locomotion represented as speed (relative to the average baseline speed) and group-level responses to repeated 26 dB re. 1 m/s^2^ stimuli (n = 27 fish). Shaded area for each trace is SEM. Speed decrease and recovery time are consistent with multiple stimulus presentations. B Histogram of path vector angles during baseline (grey) and post-stimulus (red) condition. Inset: schematic diagram showing how path vector angles are calculated. C Changes in electric field pulse evoked startle response (EFP SLC, n = 27 fish) during baseline, arrest, and recovery periods (Mean ± SEM).

**Figure S2.**
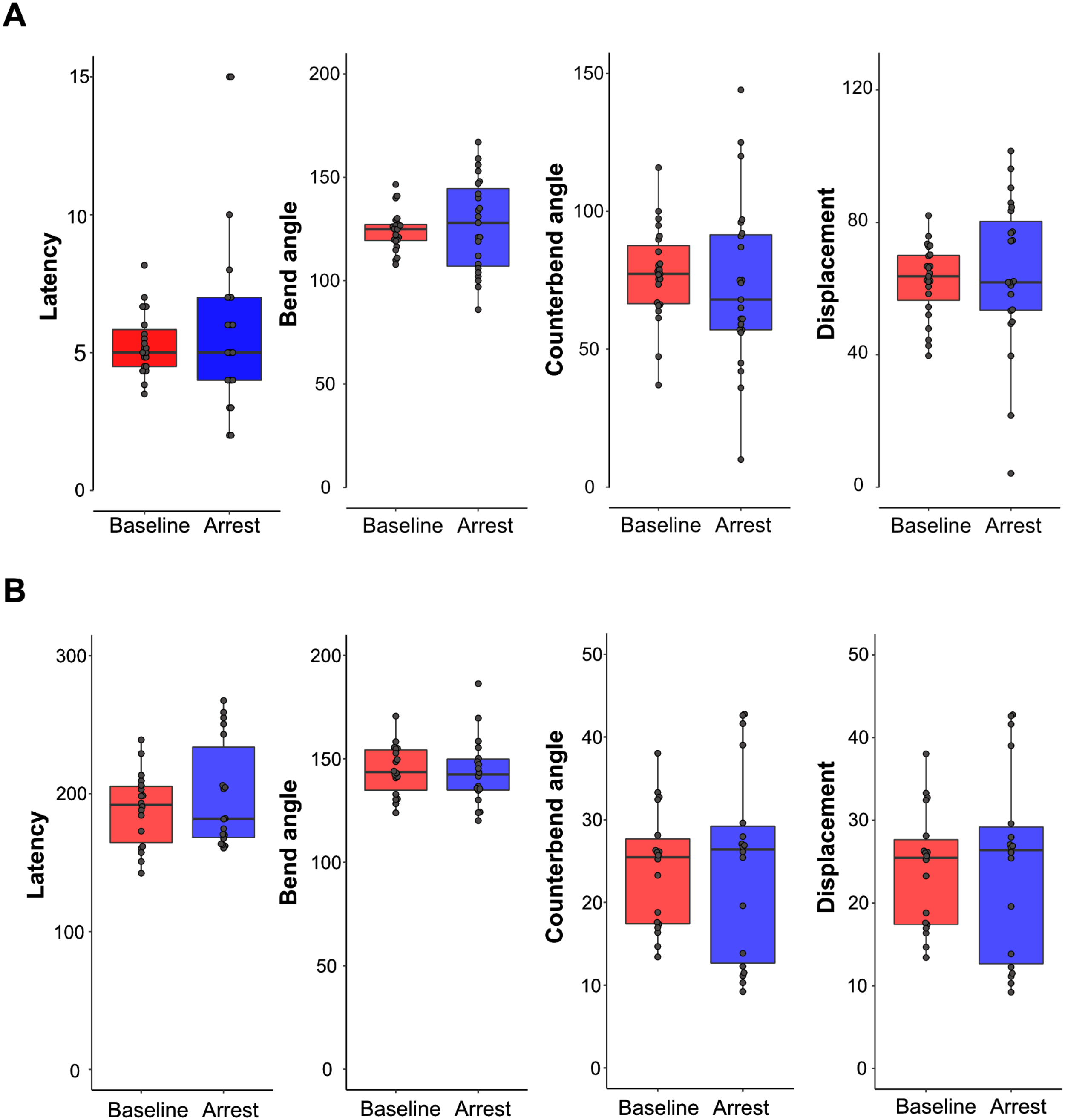
A. Kinematic analysis for short-latency startle (SLC) responses during baseline (red) and arrest (blue). No differences are observed between baseline and arrest for response latency, initial bend angle, counterbend angle, and displacement from each bout. Data points represent averaged data from each fish tested (n = 27). B. Kinematic analysis of latency, initial bend angle, counterbend angle, and displacement for dark flash O-bend responses during baseline (red) and arrest (blue).

**Figure S3.**
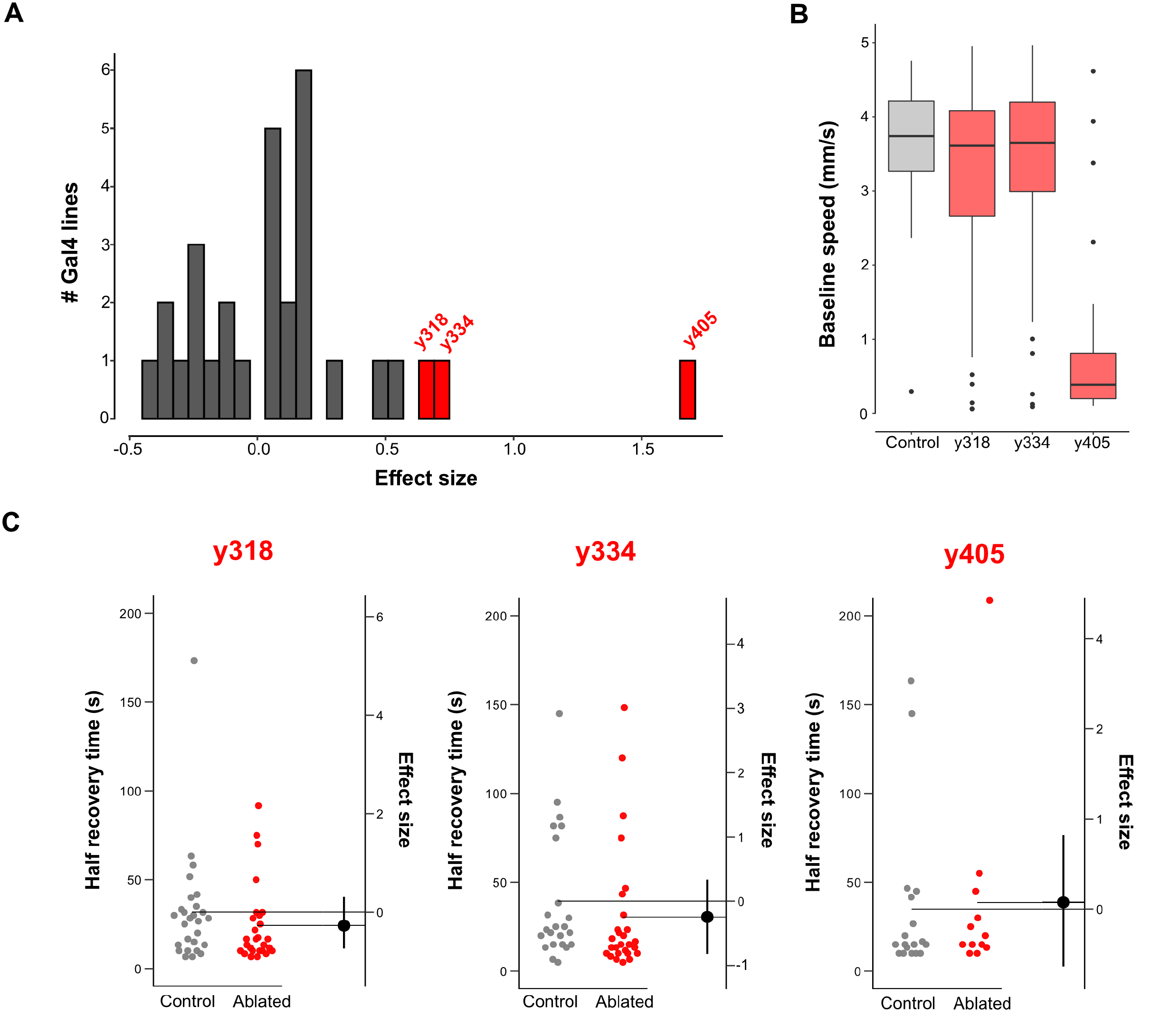
A. Histogram of changes in stimulus evoked behavioral arrest for lines used in the circuit breaking screen. Data are plotted as Cohen‘s d of differences between control and ablated fish. Gal4 ablations with high effect sizes (d > 0.6) are shown in red. B. Averaged speed (mm/s) during the pre-stimulus baseline for control fish (grey), *y318-Gal4*, *y334-Gal4*, and *y405-Gal4*. Whiskers show 95% CI. Baseline locomotion in *y405-Gal4* ablated fish is significantly reduced compared to control (p = 1.2 x 10^-14^, two-sample t-test) C. Time to half-recovery (defined as the time required to recover to 50% of pre-stimulus baseline speed) in *y318-Gal4* ablated, *y334-Gal4* ablated, and *y405-Gal4* ablated fish. Differences between control (grey) and ablated (red) fish are not statistically significant in all three ablation conditions.

**Figure S4.**
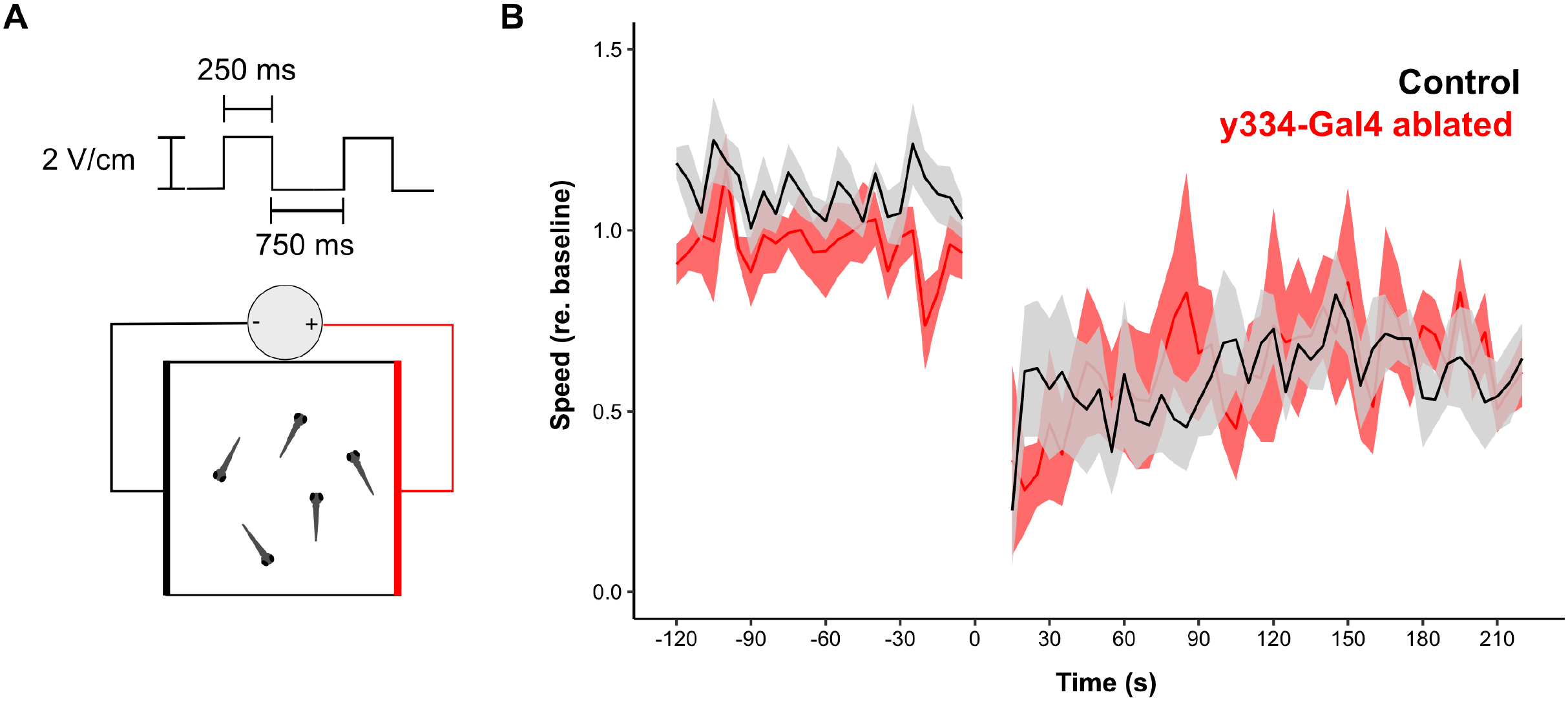
A. Schematic diagram of electrical field pulse [Top] and behavioral arena [Bottom] used for electric shock induced behavioral arrest. Groups of n = 5 fish were exposed to 2 V/cm pulsed stimuli for 30 s and total displacement was measured. B. Traces showing speed (relative to the average baseline speed) for control (black) and *y334-Gal4* ablated (red) fish. Shaded area represents SEM.

**Figure S5.**
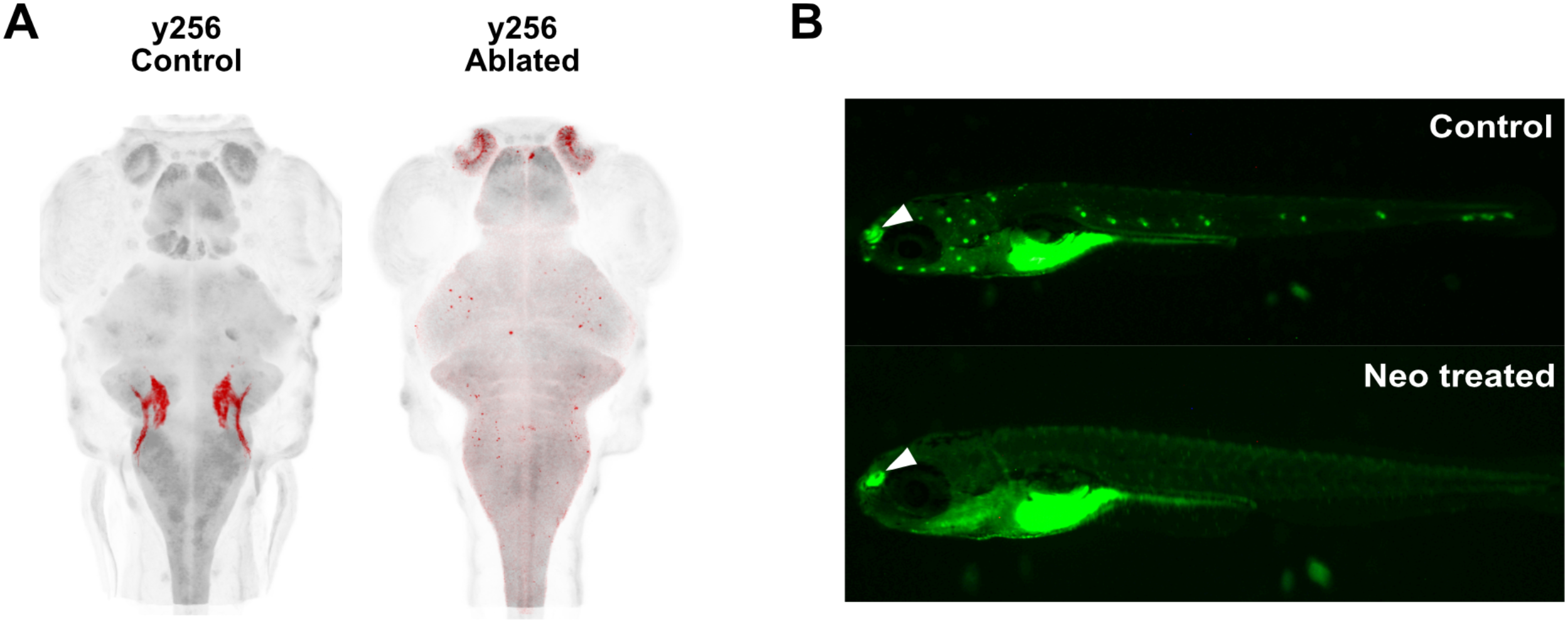
A. Example of *y256-Gal4* statoacoustic ganglion (SAG) ablation using NTR. Control (Left) and NTR ablated (Right) fish were imaged at 7 dpf after behavioral testing and registered to a reference brain. Ablated fish showed a total loss of SAG neurons. B. Example of lateral line ablation using 250 µM neomycin (Neo). DASPEI labeled neuromasts are visible in control (Top) and absent in Neo treated fish. Arrowheads show DASPEI labeling of the olfactory epithelium which is not affected by neomycin treatment.

**Figure S6.** 3D reconstructions of all traced ventral prepontine neurons from *y334-Gal4* in a model of the larval zebrafish brain (blue) with a model of RoL1 reticulospinal neurons (green). Representative examples of ipsilateral projecting (n = 7 neurons), contralateral projecting (n = 5), and hypothalamus projecting (n = 2) neurons can be seen in Figure 8B.

**Figure S7.**
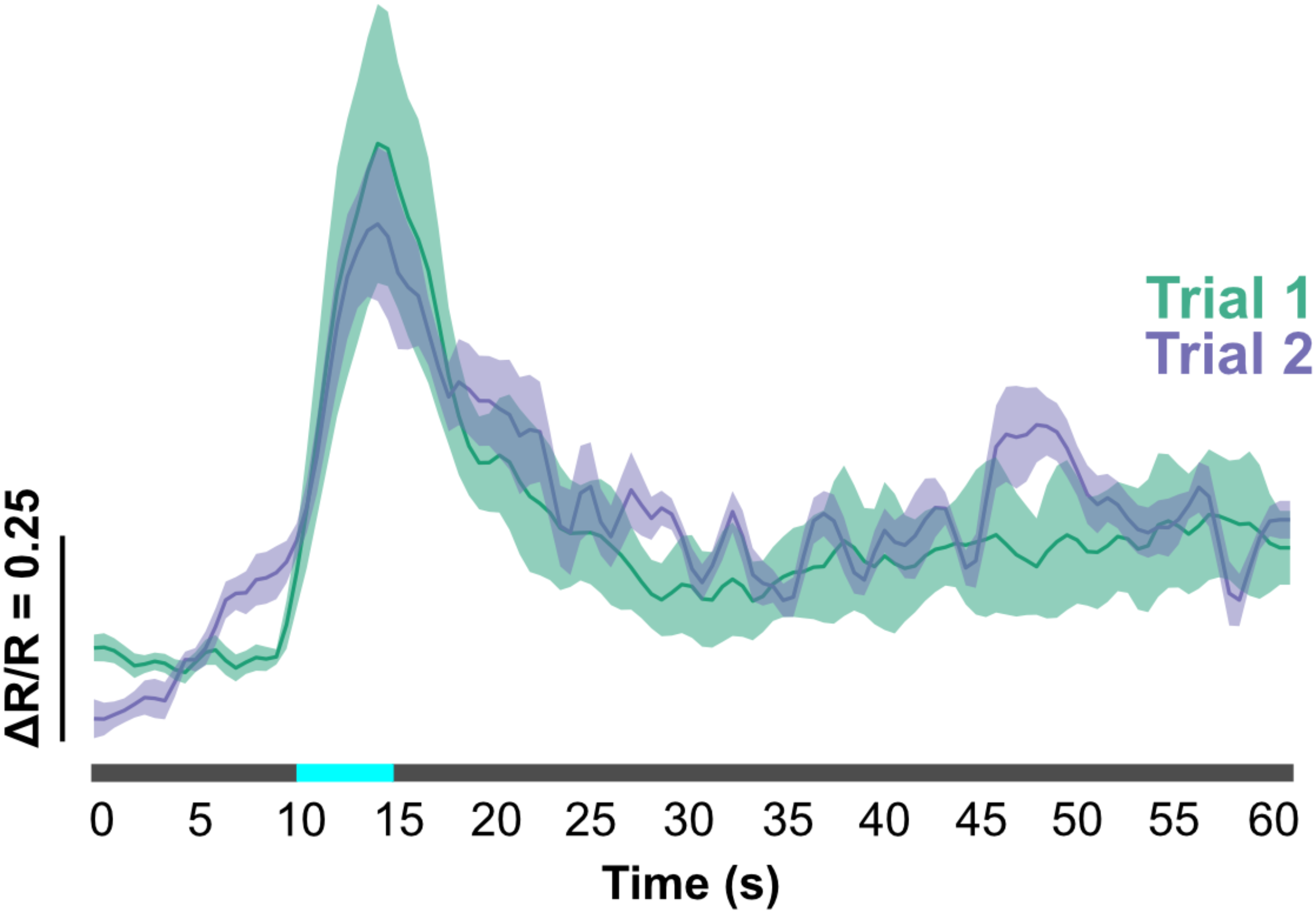
Flow-responsive vPPN responses to repeated stimuli (Average ± SEM, n = 22 cells). First stimulus presentation (Trial 1, green) and second presentation (Trial 2, purple) are strongly correlated in time and response magnitude (Pearson correlation = 0.71).

**Figure S8.**
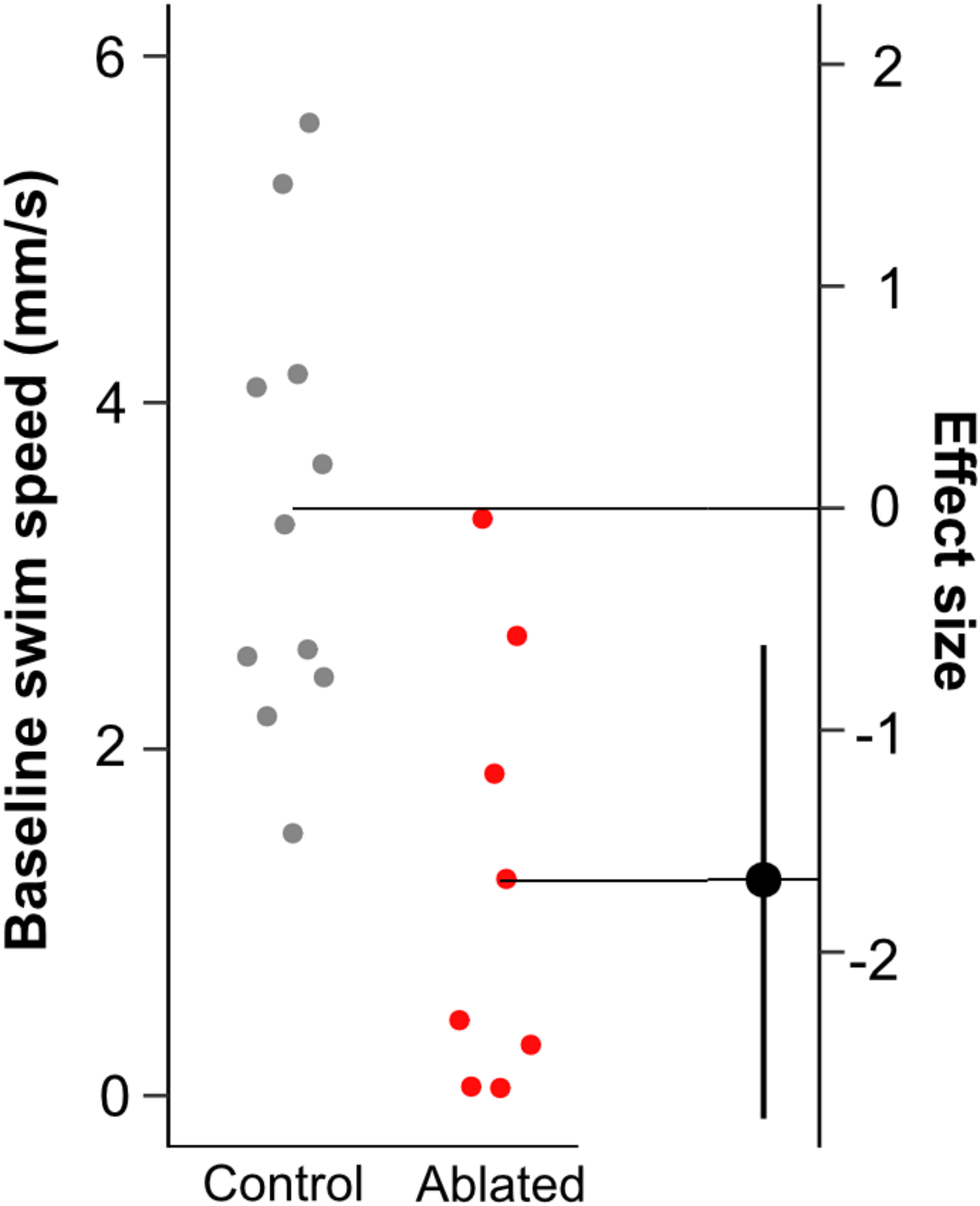
Average swim speed during the baseline period (mm/s) in RoL1 ablated neurons (red) compared to sham ablated controls (grey).

## Methods

### Zebrafish lines

All lines used in this study were maintained in a Tupfel Longfin (TL) background. Generation and description of the Gal4 lines used in this study (*y256-Gal4*, *y318-Gal4*, *y334-Gal4*, *y397-Gal4*, *y405-Gal4*) has been reported previously (Marquart et al. 2015). In addition, we received aldoca:GFF (*aldoca-Gal4*)(Takeuchi et al. 2015) as a kind gift from Masahiko Hibi (Nagoya University). Unless mentioned specifically, all Gal4 lines were maintained with and imaged with *Tg(UAS:kaede)s1999t* (Davison et al., 2007). Nitroreductase lines *Tg(UAS-E1b:BGi-epNTR-TagRFPT-oPre)y268Tg* (UAS:epNTR) and *Tg(UAS:epNTR-TagRFPT-utr.zb3)y362Tg* were used for genetic ablation experiments. Gal4-Cre intersectional ablations were conducted using *Et(REx2-SCP1:BGi-Cre-2a-Cer)y520*(Tabor et al., 2019). Other UAS lines used in this study were *Tg(UAS:EGFP-CAAX)m1230* (Fernandes et al., 2012), *Tg(14xUAS-E1b:BGi-nls-GCaMP6s.zf1-2a-nls-dsRed2.zf1)y510* (Tabor et al., 2018), *UAS:lynTagRFPT(y260)* (Yokogawa et al. 2012), *Tg(UAS-E1b:BGi-SCN5a-v2a-TagRFPT)y266* (Tabor et al. 2014), and *cUAS:PSD95.FingR-GPF-ZFC(CCR5TC)-KRAB(A)* (Son et al. 2016).

Transgenic lines used for imaging neurotransmitter identity identification were *TgBAC(slc17a6b[vglut2a]:loxP-DsRed-loxP-GFP)nns14*), *TgBAC(gad1b:LOXP-RFP-LOXP-GFP)nns26* (Satou et al., 2013), *vachta:GFP* (a kind gift from Minoru Koyama, University of Toronto), *Tg(neurod1:nsfa-EGFP)vo4Tg* (a kind gift from Katie Drerup, University of Wisconsin)(Mo and Nicolson, 2011).

The pTol1-UAS:CoChR-tdTomato (Antinucci et al., 2020) plasmid used for photoactivation experiments was a kind gift from Claire Wyart, ICM. The intracellular loop (ICL) and mutant ICL (ICLmut) constructs were generated by inserting a zebrafish codon-optimized geneblock (IDT; sequence from (Shen et al., 2009)) into UAS:dusp27-GFP (Fero et al., 2014) by replacing dusp27 with ICL using restriction enzyme digestion and ligation. Plasmids were injected with tol1 transposase and screened for fluorescence before testing.

### Behavioral experiments

All behavioral experiments were conducted at 27-28 °C. For initial behavioral arrest characterization, zebrafish larvae (6-7 dpf) were placed individually in 1 cm^2^ wells of a 3×3 arena and allowed to acclimate to ambient light for > 2 min. The arena was lit from below with a white-spectrum LED array; for experiments conducted in dark, an infrared LED was used. Sinusoidal stimuli were generated using a BNC-2110 DAQ board (National Instruments) and delivered using a Type 3810 minishaker (Bruel-Kjaer). Vibratory stimuli were calibrated so that single stimuli evoked escape responses with 83% ± 5% SEM (n = 8 fish). Behavioral responses were captured at 20 frames/sec using a µEye camera (IDS Imaging) fitted with a 40 mm macro lens (EX DG Macro, Sigma). Each trial contained a 120 sec baseline period followed by ∼15 sec vibratory stimulation and a 210 sec recovery period. Fish were imaged during the baseline and recovery period, but not during stimulation. Behavioral responses were analyzed using FLOTE software (Burgess and Granato, 2007a). Locomotion was measured as the total (x,y) distance moved within 5 sec bins and fish that displayed an average speed of < 0.5 mm/sec during the baseline period were excluded from analysis. Immobility was defined as the change in speed between baseline and the first epoch of the post-stimulus condition. This estimate of immobility is conservative because it samples a 5 sec cumulative window and because of inherent tracking variability due to instrumental noise. In cases where baseline locomotion was affected by experimental manipulation, we compared immobility in fish with baseline locomotion between 2-3 mm/s in both conditions.

Time-constant to half recovery (ΔT) was calculated as the time to recover to half of the baseline. Individuals that never recovered to half of the baseline were coded as having recovery times of 210 sec. Locomotor path analysis was conducted with locomotion data sampled at 20 Hz. After smoothing position data using a moving average, we calculated the vector angle between consecutive points using the formula 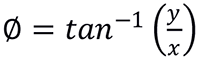

Locomotor behaviors and escape responses were imaged using the same apparatus. Responses were recorded at 1000 frames/sec using a DRS Lightning high speed camera (DEL Imaging) and analyzed using FLOTE software. Locomotor kinematics were measured in 400 ms epochs and categorized as turns or forward bouts using kinematic parameters described previously (Burgess and Granato, 2007b). Turn or forward bout initiation was calculated as the percent of epochs in which that bout was performed. For all epochs, only the first behavior was used for analysis.

Righting reflex was tested using the same apparatus used for high-speed video tracking. Groups of 5 fish were placed in a 5 cm x 5 cm arena and presented with a 15 s low frequency stimulus; control fish were not stimulated. After stimulus presentation, all fish were presented with a 1 s high amplitude, low frequency stimulus to disrupt balance. Groups of fish were tracked at 1000 frames/s for 15 sec and blinded for analysis. Righting was annotated manually by a naive observer who recorded the proportion unbalanced fish at the end of 500 ms epochs. ‘Time to balanced’ was defined as the time at which all fish in the arena showed a dorsal-up posture.

Acoustic and electric field evoked startle escape response experiments were conducted using previously published protocols (Tabor et al., 2014). Acoustic or electric pulse stimuli were delivered at 30 sec intervals to minimize habituation and categorized as short-latency or long-latency startle responses using kinematic and response latency parameters. Responsiveness was calculated as the percent of responses in each condition (baseline, SEBA, Recovery). Visual escapes using dark flash stimuli were performed using parameters published previously (Burgess & Granato 2007a). Larvae were illuminated from above with white-spectrum light for the duration of the experiment. Dark flash stimuli consisted of a total loss of illumination for 300 ms and were delivered at 60 sec intervals. Responses were characterized using kinematic parameters and responsiveness was calculated in the same manner as the SLC experiments.

### Ablations

Genetic ablations were conducted according to previously published protocols (Marquart et al., 2015; Tabor et al., 2018). Gal4 lines were chosen for the circuit breaking screen using the zebrafish brain browser (zbbrowser.com) and crossed to UAS:epNTR embryos and screened for TagRFP fluorescence at 3-4 dpf. In Gal4-Cre experiments, fish were screened for the presence of RFP in the presumed intersect.

Non-fluorescent sibling embryos were used as controls in both cases. Both groups were treated with 10 mM metronidazole (Sigma) in the absence of light for 24-48 hrs. Following ablation, larvae were washed in E3 and allowed to recover for >12 hrs before experimentation. Following behavioral experiments, a subset of ablated fish was imaged using epifluorescent microscopy to confirm full pattern ablation.

Neuromast ablations were conducted using 200 µM neomycin in E2 embryo medium according to (Harris et al., 2003). 6 dpf larvae were immersed in neomycin for 1 hr, rinsed four times in fresh E2 and allowed to recover for 3-6 hrs in E2 before testing. Following testing, ablation efficiency was confirmed using the fluorescent dye 2-4-(dimethylamino)styryl-N-ethylpyridinium-iodide (DASPEI)(Sigma). Larvae were immersed in 0.005% DASPEI in E3 for 15 min, then rinsed twice in E3. Larvae were imaged using epifluorescent microscopy with a 488 nm laser to confirm ablation. Fluorescence in the olfactory epithelium was used as a positive control of DASPEI labeling.

Multiphoton laser ablations were conducted on 4 dpf larvae raised in 300 µM N-Phenylthiourea (PTU) in E3 to suppress melanin formation. PTU was added ∼24 hpf and changed every 48 hrs. Larvae were sorted for UAS:kaede expression at 3 dpf. Larvae were anesthetized in MS222 and mounted in 2.5% low melting point agarose. Ablations were performed on a Leica TCS SPII upright confocal microscope with a MaiTai DeepSee multiphoton laser (Spectraphysics) using a 20x/1.00 NA water immersion lens. Single cells within the Gal4 pattern were identified visually using a 488 nm laser and ablated using a pulsed 800-850 nm tuned multiphoton beam. Ablations were confirmed visually using the 488 nm laser and transmitted light. Sham ablation animals were mounted in the same manner but were exposed to low laser intensity. Ablated and sham control larvae were raised in E3 until behavioral testing at 6 dpf and laser ablations were confirmed by confocal microscopy.

### Drug exposure experiments

Ethanol and GABA exposure experiments were conducted on 6 dpf fish as described above. Stock solutions of 1 mM GABA and 50% ethanol were diluted in E3 to the working concentration. Fish were immersed for 20 min in solution immediately before the experiments and controls were placed in an equal volume of E3. In the ethanol experiment, fish were immersed in 300 mM ethanol for 20 min and fish designated for the washout experiment (n=27 fish) were rinsed twice in E3 and placed into fresh E3 for an additional 20 min before testing.

### Cell capture and RNA sequencing

Cells from *y334-Gal4*;UAS:kaede and *y405-Gal4*;UAS:kaede were collected using a modified protocol from (Hempel et al., 2007). Prepontine regions from 5-6 dpf fish were dissected into Evans buffer and dissociated in 1 mg/ml neutral protease (Worthington) in Evans buffer for 30 min at room temperature with gentle shaking. The tissue fragments were rinsed 3 times with Evans buffer and triturated, and dissociated cell suspensions were plated onto a Sylgard-coated Petri dish. Kaede-expressing cells were aspirated into a glass micropipette under a fluorescent microscope. Collected cells were visually verified by plating into a fresh dish the process was repeated until pure samples of kaede-expressing cells were obtained. Groups of 6-10 cells were dispensed from the micropipette into 1 µl ice-cold 10x reaction buffer (SMART-Seq v4 kit, Takara) and flash-frozen before library preparation. Control samples were 50-100 non-prepontine cells. cDNA synthesis was performed using the SMART-Seq v4 kit according to the manufacturer‘s instructions. After cDNA purification (Beckman SPRI beads), quality control (Agilent Bioanalyzer), and quantification, samples were sequenced on an Illumina Novaseq with paired end reads. Raw read counts were co-normalized and genes with expression >1000 reads and <10 reads in control samples were filtered to remove highly expressed genes. Differential expression was calculated using Wilcoxon Rank-Sum tests and p-values were corrected for multiple comparisons using a Benjamini-Hochberg correction with a 5% FDR.

### Optogenetic activation experiments

Gal4 embryos were injected at the one cell stage with a plasmid containing UAS:CoChR with tol1 mRNA and raised in E3 under low light conditions to reduce blue light exposure. Larvae were screened for RFP fluorescence at 3 dpf and injected embryos without RFP expression were used as controls. Behavioral experiments were conducted on the same apparatus as the vibration experiments with a 470 nm LED (Prizmatix) replacing the minishaker. Experiments were done with IR illumination in darkness. Following a 10 min acclimation period, larvae were presented with a 15 sec 60 Hz pulsed stimulus to approximate the vibratory stimulus and behavior was captured and analyzed as described earlier.

### Imaging

Larvae were raised in 300 µM PTU beginning at 24 hpf. For whole brain images, 5 dpf larvae were immersed in a solution of 0.001% Lysotracker DeepRed (Invitrogen) in 1% DMSO in E3 for 12-18 hours, then rinsed in E3 twice before imaging. At 6 dpf, larvae were anesthetized in MS222 and dorsally mounted in 2.5% low melting point agarose within Lab-Tek II #1.5 cell culture chambers. In some cases, larvae expressing UAS:kaede were photoconverted using 405 nm light for 10 min before imaging. Whole and partial brain images were acquired on a Leica SPII inverted confocal microscope with a 25x/0.95 NA water immersion lens. Unless specified otherwise, images were acquired at 1 µm x 1 µm x 2 µm voxel resolution. Samples were excited using a 488 nm argon laser and a 561 nm solid state laser and detected using hybrid detectors for GFP and RFP channels and a PMT detector for far red fluorescence. Images were post-processed for dye separation using Leica Application Suite software. 3D rendering of imaged fish was conducted in Imaris 8.4.2 (Bitplane).

Individual neurons were labeled by crossing *y334-Gal4*;UAS:bloswitch fish to hsp70:B3 and raised in 300 uM PTU. Individual neurons were labeled at 3 dpf by a 20 min heat shock at 35 ℃ to activate B3 recombinase. Larvae were imaged at 6 dpf and axon projections were traced semi-automatically in Imaris using the filaments feature.

Whole brain images were acquired using 616 x 500 pixel tiles with 25 µm overlap and stitched post-hoc using custom Fiji scripts. Images were registered using previously published protocols (Marquart et al. 2015, Tabor et al. 2018). Stitched image stacks were split into individual channels and registered to a reference brain using ANTs and custom parameters. Registered images were masked with a custom mask that removed autofluorescence from skin and the eyes.

pERK/tERK labeling was conducted similar to previously published protocols (Randlett et al., 2015). Briefly, TL fish were raised in 300 uM PTU until 6 dpf. Fish were presented with arrest-inducing stimuli and fixed in 4% paraformaldehyde 2-5 min after stimulus presentation. Control fish were exposed to the vibratory stimulus, but fixed >10 min after stimulus presentation. Fixed fish were immunofluorescently labeled pERK (Phospho-pp44/42 MAP Kinase) and tERK (pp44/42 MAP Kinase) antibodies (Cell Signaling Technologies) and Alexa fluorophore conjugated secondary antibodies at 1:500 dilutions. After registration, images were masked using a vPPN mask generated from a binarized *y334-Gal4* pattern from the Zebrafish Brain Browser (zbbrowser.com). pERK/tERK ratios were calculated and averaged within the masked area for each imaged fish.

Visualization of reticulospinal neurons was conducted by backfills of *y405-Gal4* and *y334-Gal4* with UAS:kaede, UAS:LynTagRFP or UAS:caax-EGFP raised in PTU. 5 dpf larvae were anesthetized in MS222 and placed into a Sylgard coated dish. A ∼50% solution of either 3000 MW Tetramethylrhodamine biotinylated dextran or 10000 MW Alexa-488 conjugated dextran (Molecular Probes) was injected into the spinal cord immediately dorsal to the swim bladder using a PV820 pneumatic picospritzer (World Precision Instruments). Larvae were allowed to recover in Evans buffer (134 mM NaCl, 2.9 mM KCl, 2.1 mM CaCl_2_, 1.2 mM MgCl_2_, 10 mM glucose, 10 mM HEPES, pH 7.5 with NaOH) (Drapeau et al., 1999) for 24 hours and imaged as described above.

### Calcium imaging

Calcium imaging was performed on fish injected with tol1 mRNA and UAS:nls-GCamp6s-2a-nls-dsRed at the single cell stage. Larvae were raised in 300 µM PTU and screened for dsRed fluorescence at 3 dpf. At 6 dpf, larvae were mounted in 2.5% low melting point agarose in a 35 mm^2^ petri dish. The agarose surrounding the tail was cut out to allow access to water flow. Mounted larvae were placed in a custom 3D printed stage on a Leica SPII upright confocal microscope with a resonance scanner and a 20x/1.00 NA water dipping lens. Pulsed flow stimuli were delivered using a Cole-Parmer self-priming micropump controlled by a BNC-2110 DAQ board. Each stimulus set consisted of a 10 sec baseline period followed by 5 second stimulus presentation of 1 Hz water pulses and a 45 sec recovery period. In experiments with multiple stimulus presentations, stimulus sets were separated by > 120 seconds to minimize effects of consecutive stimuli. GCamp6s activity was recorded using 488 nm and 561 nm excitation at 2 Hz in single planes, which were chosen by visual approximation of the vPPN pattern. GCamp6s fluorescence was measured from image time series using custom Python scripts and the scikit-image toolbox. Nuclei position was identified from the dsRed channel using the Laplacian of Gaussian method of blob detection and was confirmed visually. Nuclear position was then registered using manual affine registration using the first 10 frames of the dsRed channel as a reference. Total fluorescence of a bounding box surrounding the detected nucleus was measured in the dsRed and GCamp6s channels and ΔR/R was normalized by dividing GCamp6s fluorescence by dsRed fluorescence. The normalized fluorescence was divided by the average baseline normalized fluorescence to obtain a final measure of fluorescence. Trials where cells drifted out of the z-plane or where the animal exhibited struggle behavior were discarded. Cells that increased fluorescence by > 3 standard deviations above the mean during the stimulus period were characterized as stimulus-responsive cells.

### Statistics

All data was analyzed using custom scripts in IDL (Harris Geospatial), R (http://www.R-project.org), and Python 3.7. All tests are two-tailed and assumed independent samples unless noted.

## References

Agetsuma, M., Aizawa, H., Aoki, T., Nakayama, R., Takahoko, M., Goto, M., Sassa, T., Amo, R., Shiraki, T., Kawakami, K., et al. (2010). The habenula is crucial for experience-dependent modification of fear responses in zebrafish. Nat. Neurosci. 13, 1354–1356.

Aizenman, C.D., and Linden, D.J. (1999). Regulation of the rebound depolarization and spontaneous firing patterns of deep nuclear neurons in slices of rat cerebellum. J. Neurophysiol. 82, 1697–1709.

Alderman, S.L., and Bernier, N.J. (2009). Ontogeny of the corticotropin-releasing factor system in zebrafish. Gen. Comp. Endocrinol. 164, 61–69.

Anderson, D.J., and Adolphs, R. (2014). A framework for studying emotions across species. Cell 157, 187–200.

Antinucci, P., Dumitrescu, A., Deleuze, C., Morley, H.J., Leung, K., Hagley, T., Kubo, F., Baier, H., Bianco, I.H., and Wyart, C. (2020). A calibrated optogenetic toolbox of stable zebrafish opsin lines. Elife 9.

Assad, N., Luz, W.L., Santos-Silva, M., Carvalho, T., Moraes, S., Picanço-Diniz, D.L.W., Bahia, C.P., Oliveira Batista, E. de J., da Conceição Passos, A., Oliveira, K.R.H.M., et al. (2020). Acute Restraint Stress Evokes Anxiety-Like Behavior Mediated by Telencephalic Inactivation and GabAergic Dysfunction in Zebrafish Brains. Sci. Rep. 10, 5551.

Bae, Y.-K., Kani, S., Shimizu, T., Tanabe, K., Nojima, H., Kimura, Y., Higashijima, S.-I., and Hibi, M. (2009). Anatomy of zebrafish cerebellum and screen for mutations affecting its development. Dev. Biol. 330, 406–426.

Bagnall, M.W., and McLean, D.L. (2014). Modular organization of axial microcircuits in zebrafish. Science 343, 197–200.

Barik, A., Thompson, J.H., Seltzer, M., Ghitani, N., and Chesler, A.T. (2018). A Brainstem-Spinal Circuit Controlling Nocifensive Behavior. Neuron 100, 1491–1503.e3.

Barrios, J.P., Wang, W.-C., England, R., Reifenberg, E., and Douglass, A.D. (2020). Hypothalamic dopamine neurons control sensorimotor behavior by modulating brainstem premotor nuclei.

Bergeron, S.A., Hannan, M.C., Codore, H., Fero, K., Li, G.H., Moak, Z., Yokogawa, T., and Burgess, H.A. (2012). Brain selective transgene expression in zebrafish using an NRSE derived motif. Front. Neural Circuits 6, 110.

Bergeron, S.A., Carrier, N., Li, G.H., Ahn, S., and Burgess, H.A. (2015). Gsx1 expression defines neurons required for prepulse inhibition. Mol. Psychiatry 20, 974–985.

Bowen, A.J., Chen, J.Y., Huang, Y.W., Baertsch, N.A., Park, S., and Palmiter, R.D. (2020). Dissociable control of unconditioned responses and associative fear learning by parabrachial CGRP neurons. Elife 9.

Burgess, H.A., and Granato, M. (2007a). Modulation of locomotor activity in larval zebrafish during light adaptation. J. Exp. Biol. 210, 2526–2539.

Burgess, H.A., and Granato, M. (2007b). Sensorimotor gating in larval zebrafish. J. Neurosci. 27, 4984– 4994.

Cachat, J., Stewart, A., Grossman, L., Gaikwad, S., Kadri, F., Chung, K.M., Wu, N., Wong, K., Roy, S., Suciu, C., et al. (2010). Measuring behavioral and endocrine responses to novelty stress in adult zebrafish. Nat. Protoc. 5, 1786–1799.

Carli, G. (1968). Depression of somatic reflexes during rabbit hypnosis. Brain Res. 11, 453–456.

Chandrasekar, G., Lauter, G., and Hauptmann, G. (2007). Distribution of corticotropin-releasing hormone in the developing zebrafish brain. J. Comp. Neurol. 505, 337–351.

Chiang, M.C., Bowen, A., Schier, L.A., Tupone, D., Uddin, O., and Heinricher, M.M. (2019). Parabrachial Complex: A Hub for Pain and Aversion. J. Neurosci. 39, 8225–8230.

Crawford, F.T. (1977). Induction and Duration of Tonic Immobility. Psychol. Rec. 27, 89–107.

Dai, Y., Iwata, K., Fukuoka, T., Kondo, E., Tokunaga, A., Yamanaka, H., Tachibana, T., Liu, Y., and Noguchi, K. (2002). Phosphorylation of extracellular signal-regulated kinase in primary afferent neurons by noxious stimuli and its involvement in peripheral sensitization. J. Neurosci. 22, 7737–7745.

Davison, J.M., Akitake, C.M., Goll, M.G., Rhee, J.M., Gosse, N., Baier, H., Halpern, M.E., Leach, S.D., and Parsons, M.J. (2007). Transactivation from Gal4-VP16 transgenic insertions for tissue-specific cell labeling and ablation in zebrafish. Dev. Biol. 304, 811–824.

Dean, P., Redgrave, P., and Westby, G.W. (1989). Event or emergency? Two response systems in the mammalian superior colliculus. Trends Neurosci. 12, 137–147.

Dohaku, R., Yamaguchi, M., Yamamoto, N., Shimizu, T., Osakada, F., and Hibi, M. (2019). Tracing of Afferent Connections in the Zebrafish Cerebellum Using Recombinant Rabies Virus. Front. Neural Circuits 13, 30.

Drapeau, P., Ali, D.W., Buss, R.R., and Saint-Amant, L. (1999). In vivo recording from identifiable neurons of the locomotor network in the developing zebrafish. J. Neurosci. Methods 88, 1–13.

Duboué, E.R., Hong, E., Eldred, K.C., and Halpern, M.E. (2017). Left Habenular Activity Attenuates Fear Responses in Larval Zebrafish. Curr. Biol. 27, 2154–2162.e3.

Edson, P.H., and Gallup, G.G. (1972). Tonic immobility as a fear response in lizards Anolis carolinensis. Psychon. Sci. 26, 27–28.

Ekker, M., Wegner, J., Akimenko, M.A., and Westerfield, M. (1992). Coordinate embryonic expression of three zebrafish engrailed genes. Development 116, 1001–1010.

Esposito, G., Yoshida, S., Ohnishi, R., Tsuneoka, Y., Rostagno, M.D.C., Yokota, S., Okabe, S., Kamiya, K., Hoshino, M., Shimizu, M., et al. (2013). Infant calming responses during maternal carrying in humans and mice. Curr. Biol. 23, 739–745.

Fanselow, M.S. (1994). Neural organization of the defensive behavior system responsible for fear. Psychon. Bull. Rev. 1, 429–438.

Favre-Bulle, I.A., Stilgoe, A.B., Rubinsztein-Dunlop, H., and Scott, E.K. (2017). Optical trapping of otoliths drives vestibular behaviours in larval zebrafish. Nat. Commun. 8, 630.

Fernandes, A.M., Fero, K., Arrenberg, A.B., Bergeron, S.A., Driever, W., and Burgess, H.A. (2012). Deep brain photoreceptors control light-seeking behavior in zebrafish larvae. Curr. Biol. 22, 2042–2047.

Fero, K., Yokogawa, T., and Burgess, H.A. (2011). The Behavioral Repertoire of Larval Zebrafish. In Zebrafish Models in Neurobehavioral Research, A.V. Kalueff, and J.M. Cachat, eds. (Totowa, NJ: Humana Press), pp. 249–291.

Fero, K., Bergeron, S.A., Horstick, E.J., Codore, H., Li, G.H., Ono, F., Dowling, J.J., and Burgess, H.A. (2014). Impaired embryonic motility in dusp27 mutants reveals a developmental defect in myofibril structure. Dis. Model. Mech. 7, 289–298.

Förster, D., Dal Maschio, M., Laurell, E., and Baier, H. (2017). An optogenetic toolbox for unbiased discovery of functionally connected cells in neural circuits. Nat. Commun. 8, 116.

Fulwiler, C.E., and Saper, C.B. (1984). Subnuclear organization of the efferent connections of the parabrachial nucleus in the rat. Brain Res. 319, 229–259.

Gahtan, E., and O‘Malley, D.M. (2003). Visually guided injection of identified reticulospinal neurons in zebrafish: a survey of spinal arborization patterns. J. Comp. Neurol. 459, 186–200.

Gallup, G.G., Jr. (1977). Tonic immobility: The role of fear and predation. Psychol. Rec. 27, 41–61.

Gallup, G.G., and Rager, D.R. (1996). Tonic Immobility as a Model of Extreme States of Behavioral Inhibition. In Motor Activity and Movement Disorders: Research Issues and Applications, P.R. Sanberg, K.-P. Ossenkopp, and M. Kavaliers, eds. (Totowa, NJ: Humana Press), pp. 57–80.

Gibson, W.T., Gonzalez, C.R., Fernandez, C., Ramasamy, L., Tabachnik, T., Du, R.R., Felsen, P.D., Maire, M.R., Perona, P., and Anderson, D.J. (2015). Behavioral responses to a repetitive visual threat stimulus express a persistent state of defensive arousal in Drosophila. Curr. Biol. 25, 1401–1415.

Han, S., Soleiman, M.T., Soden, M.E., Zweifel, L.S., and Palmiter, R.D. (2015). Elucidating an Affective Pain Circuit that Creates a Threat Memory. Cell 162, 363–374.

Harris, J. a., Cheng, A.G., Cunningham, L.L., MacDonald, G., Raible, D.W., and Rubel, E.W. (2003). Neomycin-induced hair cell death and rapid regeneration in the lateral line of zebrafish (Danio rerio). J. Assoc. Res. Otolaryngol. 4, 219–234.

Hempel, C.M., Sugino, K., and Nelson, S.B. (2007). A manual method for the purification of fluorescently labeled neurons from the mammalian brain. Nat. Protoc. 2, 2924–2929.

Henningsen, A.D. (1994). Tonic immobility in 12 elasmobranchs: Use as an aid in captive husbandry. Zoo Biol. 13, 325–332.

Henriques, P.M., Rahman, N., Jackson, S.E., and Bianco, I.H. (2019). Nucleus Isthmi Is Required to Sustain Target Pursuit during Visually Guided Prey-Catching. Curr. Biol. 0.

Horstick, E.J., Jordan, D.C., Bergeron, S.A., Tabor, K.M., Serpe, M., Feldman, B., and Burgess, H.A. (2015). Increased functional protein expression using nucleotide sequence features enriched in highly expressed genes in zebrafish. Nucleic Acids Res. 43, e48.

Hsieh, J.-Y., Ulrich, B., Issa, F.A., Wan, J., and Papazian, D.M. (2014). Rapid development of Purkinje cell excitability, functional cerebellar circuit, and afferent sensory input to cerebellum in zebrafish. Front. Neural Circuits 8, 147.

Humphreys, R.K., and Ruxton, G.D. (2018). A review of thanatosis (death feigning) as an anti-predator behaviour. Behav. Ecol. Sociobiol. 72, 22.

Ishikawa, T., Shimuta, M., and Häusser, M. (2015). Multimodal sensory integration in single cerebellar granule cells in vivo. Elife 4.

Jänicke, B., and Coper, H. (1996). Tests in Rodents for Assessing Sensorimotor Performance During Aging. In Advances in Psychology, A.-M. Ferrandez, and N. Teasdale, eds. (North-Holland), pp. 201– 233.

Jesuthasan, S., Krishnan, S., Cheng, R.-K., and Mathuru, A. (2020). Neural correlates of state transitions elicited by a chemosensory danger cue. Prog. Neuropsychopharmacol. Biol. Psychiatry 110110.

Kalaf, J., Vilete, L.M.P., Volchan, E., Fiszman, A., Coutinho, E.S.F., Andreoli, S.B., Quintana, M.I., de Jesus Mari, J., and Figueira, I. (2015). Peritraumatic tonic immobility in a large representative sample of the general population: association with posttraumatic stress disorder and female gender. Compr. Psychiatry 60, 68–72.

Kinkhabwala, A., Riley, M., Koyama, M., Monen, J., Satou, C., Kimura, Y., Higashijima, S.-I., and Fetcho, J. (2011). A structural and functional ground plan for neurons in the hindbrain of zebrafish. Proc. Natl. Acad. Sci. U. S. A. 108, 1164–1169.

Klemm, W.R. (1976). Identity of sensory and motor systems that are critical to the immobility reflex (―animal hypnosis‖). J. Neurosci. Res. 2, 57–69.

Klemm, W.R. (2001). Behavioral arrest: in search of the neural control system. Prog. Neurobiol. 65, 453– 471.

Koutsikou, S., Crook, J.J., Earl, E.V., Leith, J.L., Watson, T.C., Lumb, B.M., and Apps, R. (2014). Neural substrates underlying fear-evoked freezing: the periaqueductal grey-cerebellar link. J. Physiol. 592, 2197– 2213.

Kozlowska, K., Walker, P., McLean, L., and Carrive, P. (2015). Fear and the Defense Cascade: Clinical Implications and Management. Harv. Rev. Psychiatry 23, 263–287.

LeDoux, J.E. (2000). Emotion circuits in the brain. Annu. Rev. Neurosci. 23, 155–184.

LeDoux, J., and Daw, N.D. (2018). Surviving threats: neural circuit and computational implications of a new taxonomy of defensive behaviour. Nat. Rev. Neurosci. 19, 269–282.

Lefebvre, L., and Sabourin, M. (1977). Effects of spaced and massed repeated elicitation on tonic immobility in the goldfish (Carassius auratus). Behav. Biol. 21, 300–305.

Liang, F., Xiong, X.R., Zingg, B., Ji, X.-Y., Zhang, L.I., and Tao, H.W. (2015). Sensory Cortical Control of a Visually Induced Arrest Behavior via Corticotectal Projections. Neuron 86, 755–767.

Lovett-Barron, M., Chen, R., Bradbury, S., Andalman, A.S., Wagle, M., Guo, S., and Deisseroth, K. (2020). Multiple convergent hypothalamus–brainstem circuits drive defensive behavior. Nat. Neurosci.

Marquart, G.D., Tabor, K.M., Brown, M., Strykowski, J.L., Varshney, G.K., LaFave, M.C., Mueller, T., Burgess, S.M., Higashijima, S.-I., and Burgess, H.A. (2015). A 3D Searchable Database of Transgenic Zebrafish Gal4 and Cre Lines for Functional Neuroanatomy Studies. Front. Neural Circuits 9, 78.

Marques, J.C., Lackner, S., Félix, R., and Orger, M.B. (2018). Structure of the Zebrafish Locomotor Repertoire Revealed with Unsupervised Behavioral Clustering. Curr. Biol. 0.

Maruska, K.P., and Tricas, T.C. (2009). Central projections of octavolateralis nerves in the brain of a soniferous damselfish (Abudefduf abdominalis). J. Comp. Neurol. 512, 628–650.

Marx, B.P., Forsyth, J.P., Gallup, G.G., Fusé, T., and Lexington, J.M. (2008). Tonic Immobility as an Evolved Predator Defense: Implications for Sexual Assault Survivors. Clin Psychol Sci & Pract 15, 74– 90.

Matsui, H., Namikawa, K., Babaryka, A., and Köster, R.W. (2014). Functional regionalization of the teleost cerebellum analyzed in vivo. Proc. Natl. Acad. Sci. U. S. A. 111, 11846–11851.

Maximino, C., do Carmo Silva, R.X., Dos Santos Campos, K., de Oliveira, J.S., Rocha, S.P., Pyterson, M.P., Dos Santos Souza, D.P., Feitosa, L.M., Ikeda, S.R., Pimentel, A.F.N., et al. (2019). Sensory ecology of ostariophysan alarm substances. J. Fish Biol. 95, 274–286.

McBride, R.L., and Klemm, W.R. (1969). Mechanisms of the immobility reflex.

McCormick, C.A., Gallagher, S., Cantu-Hertzler, E., and Woodrick, S. (2016). Mechanosensory Lateral Line Nerve Projections to Auditory Neurons in the Dorsal Descending Octaval Nucleus in the Goldfish, Carassius auratus. Brain Behav. Evol. 88, 68–80.

Menescal-de-Oliveira, L., and Hoffmann, A. (1993). The parabrachial region as a possible region modulating simultaneously pain and tonic immobility. Behav. Brain Res. 56, 127–132.

Mo, W., and Nicolson, T. (2011). Both pre- and postsynaptic activity of Nsf prevents degeneration of hair-cell synapses. PLoS One 6, e27146.

New, J.G., and Northcutt, R.G. (1984). Central projections of the lateral line nerves in the shovelnose sturgeon. J. Comp. Neurol. 225, 129–140.

Orger, M.B., Kampff, A.R., Severi, K.E., Bollmann, J.H., and Engert, F. (2008). Control of visually guided behavior by distinct populations of spinal projection neurons. Nat. Neurosci. 11, 327–333.

Palmiter, R.D. (2018). The Parabrachial Nucleus: CGRP Neurons Function as a General Alarm. Trends Neurosci. 41, 280–293.

Perrins, R., Walford, A., and Roberts, A. (2002). Sensory Activation and Role of Inhibitory Reticulospinal Neurons that Stop Swimming in Hatchling Frog Tadpoles. J. Neurosci. 22, 4229–4240.

Randlett, O., Wee, C.L., Naumann, E. a., Nnaemeka, O., Schoppik, D., Fitzgerald, J.E., Portugues, R., Lacoste, A.M.B., Riegler, C., Engert, F., et al. (2015). Whole-brain activity mapping onto a zebrafish brain atlas. Nat. Methods 12, 1–12.

Roelofs, K. (2017). Freeze for action: neurobiological mechanisms in animal and human freezing. Philos. Trans. R. Soc. Lond. B Biol. Sci. 372.

Rogers, S.M., and Simpson, S.J. (2014). Thanatosis. Curr. Biol. 24, R1031–3.

Roseberry, T., and Kreitzer, A. (2017). Neural circuitry for behavioural arrest. Philos. Trans. R. Soc. Lond. B Biol. Sci. 372.

Ryan, S.J., Ehrlich, D.E., Jasnow, A.M., Daftary, S., Madsen, T.E., and Rainnie, D.G. (2012). Spike-timing precision and neuronal synchrony are enhanced by an interaction between synaptic inhibition and membrane oscillations in the amygdala. PLoS One 7, e35320.

Saab, C.Y., and Willis, W.D. (2003). The cerebellum: organization, functions and its role in nociception. Brain Res. Brain Res. Rev. 42, 85–95.

Satou, C., Kimura, Y., Hirata, H., Suster, M.L., Kawakami, K., and Higashijima, S.-I. (2013). Transgenic tools to characterize neuronal properties of discrete populations of zebrafish neurons. Development 140, 3927–3931.

Sawtell, N.B. (2010). Multimodal integration in granule cells as a basis for associative plasticity and sensory prediction in a cerebellum-like circuit. Neuron 66, 573–584.

Shen, W., Da Silva, J.S., He, H., and Cline, H.T. (2009). Type A GABA-receptor-dependent synaptic transmission sculpts dendritic arbor structure in Xenopus tadpoles in vivo. J. Neurosci. 29, 5032–5043.

Shen, W., McKeown, C.R., Demas, J.A., and Cline, H.T. (2011). Inhibition to excitation ratio regulates visual system responses and behavior in vivo. J. Neurophysiol. 106, 2285–2302.

Sherman, J.E., and Kalin, N.H. (1988). ICV-CRH alters stress-induced freezing behavior without affecting pain sensitivity. Pharmacology Biochemistry and Behavior 30, 801–807.

Sisneros, J.A., Tricas, T.C., and Luer, C.A. (1998). Response properties and biological function of the skate electrosensory system during ontogeny. J. Comp. Physiol. A 183, 87–99.

Sohal, V.S., Pangratz-Fuehrer, S., Rudolph, U., and Huguenard, J.R. (2006). Intrinsic and synaptic dynamics interact to generate emergent patterns of rhythmic bursting in thalamocortical neurons. J. Neurosci. 26, 4247–4255.

Son, J.-H., Keefe, M.D., Stevenson, T.J., Barrios, J.P., Anjewierden, S., Newton, J.B., Douglass, A.D., and Bonkowsky, J.L. (2016). Transgenic FingRs for Live Mapping of Synaptic Dynamics in Genetically-Defined Neurons. Sci. Rep. 6, 18734.

Speedie, N., and Gerlai, R. (2008). Alarm substance induced behavioral responses in zebrafish (Danio rerio). Behav. Brain Res. 188, 168–177.

Spinieli, R.L., and Leite-Panissi, C.R.A. (2018). Similar effect of CRF1 and CRF2 receptor in the basolateral or central nuclei of the amygdala on tonic immobility behavior. Brain Res. Bull. 137, 187– 196.

Supple, W.F., Jr, Cranney, J., and Leaton, R.N. (1988). Effects of lesions of the cerebellar vermis on VMH lesion-induced hyperdefensiveness, spontaneous mouse killing, and freezing in rats. Physiol. Behav. 42, 145–153.

Tabor, K.M., Bergeron, S. a., Horstick, E.J., Jordan, D.C., Aho, V., Porkka-Heiskanen, T., Haspel, G., and Burgess, H. a. (2014). Direct activation of the Mauthner cell by electric field pulses drives ultrarapid escape responses. J. Neurophysiol. 112, 834–844.

Tabor, K.M., Smith, T.S., Brown, M., Bergeron, S.A., Briggman, K.L., and Burgess, H.A. (2018). Presynaptic Inhibition Selectively Gates Auditory Transmission to the Brainstem Startle Circuit. Curr. Biol. 28, 2527–2535.e8.

Tabor, K.M., Marquart, G.D., Hurt, C., Smith, T.S., Geoca, A.K., Bhandiwad, A.A., Subedi, A., Sinclair, J.L., Rose, H.M., Polys, N.F., et al. (2019). Brain-wide cellular resolution imaging of Cre transgenic zebrafish lines for functional circuit-mapping. Elife 8.

Takeuchi, M., Matsuda, K., Yamaguchi, S., Asakawa, K., Miyasaka, N., Lal, P., Yoshihara, Y., Koga, A., Kawakami, K., Shimizu, T., et al. (2015). Establishment of Gal4 transgenic zebrafish lines for analysis of development of cerebellar neural circuitry. Dev. Biol. 397, 1–17.

Tovote, P., Esposito, M.S., Botta, P., Chaudun, F., Fadok, J.P., Markovic, M., Wolff, S.B.E., Ramakrishnan, C., Fenno, L., Deisseroth, K., et al. (2016). Midbrain circuits for defensive behaviour. Nature 534, 206–212.

Tremere, L.A., Pinaud, R., Irwin, R.P., and Allen, C.N. (2008). Postinhibitory rebound spikes are modulated by the history of membrane hyperpolarization in the SCN. Eur. J. Neurosci. 28, 1127–1135.

Vaaga, C.E., Brown, S.T., and Raman, I.M. (2020). Cerebellar modulation of synaptic input to freezing-related neurons in the periaqueductal gray. Elife 9.

Volchan, E., Rocha-Rego, V., Bastos, A.F., Oliveira, J.M., Franklin, C., Gleiser, S., Berger, W., Souza, G.G.L., Oliveira, L., David, I.A., et al. (2017). Immobility reactions under threat: A contribution to human defensive cascade and PTSD. Neurosci. Biobehav. Rev. 76, 29–38.

Volkmann, K., Rieger, S., Babaryka, A., and Köster, R.W. (2008). The zebrafish cerebellar rhombic lip is spatially patterned in producing granule cell populations of different functional compartments. Dev. Biol. 313, 167–180.

Watson, C., Bartholomaeus, C., and Puelles, L. (2019). Time for Radical Changes in Brain Stem Nomenclature-Applying the Lessons From Developmental Gene Patterns. Front. Neuroanat. 13, 10.

Witter, L., Canto, C.B., Hoogland, T.M., de Gruijl, J.R., and De Zeeuw, C.I. (2013). Strength and timing of motor responses mediated by rebound firing in the cerebellar nuclei after Purkinje cell activation. Front. Neural Circuits 7, 133.

Yeomans, J.S., and Frankland, P.W. (1995). The acoustic startle reflex: neurons and connections. Brain Res. Brain Res. Rev. 21, 301–314.

Yokogawa, T., Hannan, M.C., and Burgess, H. a. (2012). The Dorsal Raphe Modulates Sensory Responsiveness during Arousal in Zebrafish. J. Neurosci. 32, 15205–15215.

Yoshida, M. (2021). Immobility Behaviors in Fish: A Comparison with Other Vertebrates. In Death-Feigning in Insects: Mechanism and Function of Tonic Immobility, M. Sakai, ed. (Singapore: Springer Singapore), pp. 159–178.

Zacarias, R., Namiki, S., Card, G.M., Vasconcelos, M.L., and Moita, M.A. (2018). Speed dependent descending control of freezing behavior in Drosophila melanogaster. Nat. Commun. 9, 3697.

Zheng, N., and Raman, I.M. (2009). Ca currents activated by spontaneous firing and synaptic disinhibition in neurons of the cerebellar nuclei. J. Neurosci. 29, 9826–9838.

